# Thermodynamic and kinetic analysis of the LAO binding protein and its isolated domains reveal non-additivity in stability, folding, and function

**DOI:** 10.1101/2023.03.18.532921

**Authors:** Renan Vergara, Tania Berrocal, Eva Isela Juárez Mejía, Sergio Romero-Romero, Isabel Velázquez-López, Nancy O. Pulido, Haven A. López Sanchez, Daniel-Adriano Silva, Miguel Costas, Adela Rodríguez-Romero, Rogelio Rodríguez-Sotres, Alejandro Sosa-Peinado, D. Alejandro Fernández-Velasco

## Abstract

Substrate-binding proteins (SBP) are used by organisms from the three domains of life for transport and signaling. SBPs are composed of two domains that collectively trap ligands with high affinity and selectivity. To explore the role of the domains and the integrity of the hinge region between them in the function and conformation of SBPs, here we describe the ligand binding, conformational stability, and folding kinetics of the Lysine Arginine Ornithine binding protein (LAO) from *Salmonella thiphimurium* and constructs corresponding to its two independent domains. LAO is a class II SBP formed by a continuous and a discontinuous domain. Contrary to the expected behavior based on their connectivity, the discontinuous domain shows a stable native-like structure that binds L-arginine with moderate affinity, whereas the continuous domain is barely stable and shows no detectable ligand binding. Regarding folding kinetics, studies of the entire protein revealed the presence of at least two intermediates. While the unfolding and refolding of the continuous domain exhibited only a single intermediate and simpler and faster kinetics than LAO, the folding mechanism of the discontinuous domain was complex and involved multiple intermediates. These findings suggest that in the complete protein the continuous domain nucleates folding and that its presence funnels the folding of the discontinuous domain avoiding nonproductive interactions. The strong dependence of the function, stability, and folding pathway of the lobes on their covalent association, is most likely the result of the coevolution of both domains as a single unit.

## Introduction

Research on protein folding over the last 50 years [1] has been critical for the progress achieved in important areas of protein science, such as a mechanistic understanding of protein folding diseases [2–7] and the development of de novo protein design [8–12]. Most of our knowledge about protein folding comes from studies performed with small (< 200 aa), single-domain polypeptide chains, whereas proteomes are mainly composed of larger proteins commonly containing more than one domain [13–16]. Domain-domain interactions are also essential for function when binding and/or catalytic residues are located at or near domain-domain interphases [17]. Folding studies of whole multidomain proteins compared to those of their constitutive domains exhibit a diversified picture. In some cases, domain association impacts stability and folding kinetics are influenced by domain association [16,18–22] whereas in other cases additivity is observed [23,24]. Cooperativity between domains has been proposed to depend on the packing density and size of the domain interaction, however, only a reduced set of proteins has been studied so far, and exceptions have been found [25]. Among the obstacles encountered in the thermodynamic study of multidomain proteins, the irreversibility and the kinetic control observed during their folding transition induced by temperature or chemical agents are the hardest to overcome. As an alternative, the role of domain interactions in the folding/unfolding of very complex multidomain proteins such as chaperones has been studied using force as perturbant by single-molecule force spectroscopy [26–29].

The substrate-binding proteins (SBPs) are a family of bidomain proteins that generally show high reversibility in their temperature-, force- and chaotropic-induced unfolding transitions [30–34]. SBPs present a common general structure composed of two Rossman-fold domains linked by a hinge region. Ligand binding induces a conformational change from an open to closed state where the two domains approach each other and shield the ligand from the solvent [35–37]. SBPs are associated with membrane complexes implicated in transport and signaling, such as ABC transporters (where SBP were initially identified), two-component regulatory systems, ionic channels, and G protein-coupled receptors [35,36].

Periplasmic binding proteins (PBPs) are a category of SBDs involved in transporting several substrates into the cytoplasm of gram-negative bacteria [38–41]. The lysine-arginine-ornithine binding protein (LAO) is a type II PBP composed of one discontinuous and one continuous domain, referred to as domain A and B, respectively (Figure 1) [42]. Ligand binding to LAO exhibits high affinity and exquisite selectivity due to the interplay of both water-mediated and direct protein-ligand contacts [43]. The structural and functional role of domain-domain interactions has been previously described for HisJ, a closely related PBP, where the domains were expressed and characterized independently [44]. In order to elucidate the contribution of both domains and the integrity of the hinge region connecting them in the function and conformation of LAO, we have studied the folding, structure and ligand binding of this PBP, as well as constructs corresponding to its two independent domains.

**Figure 1.**
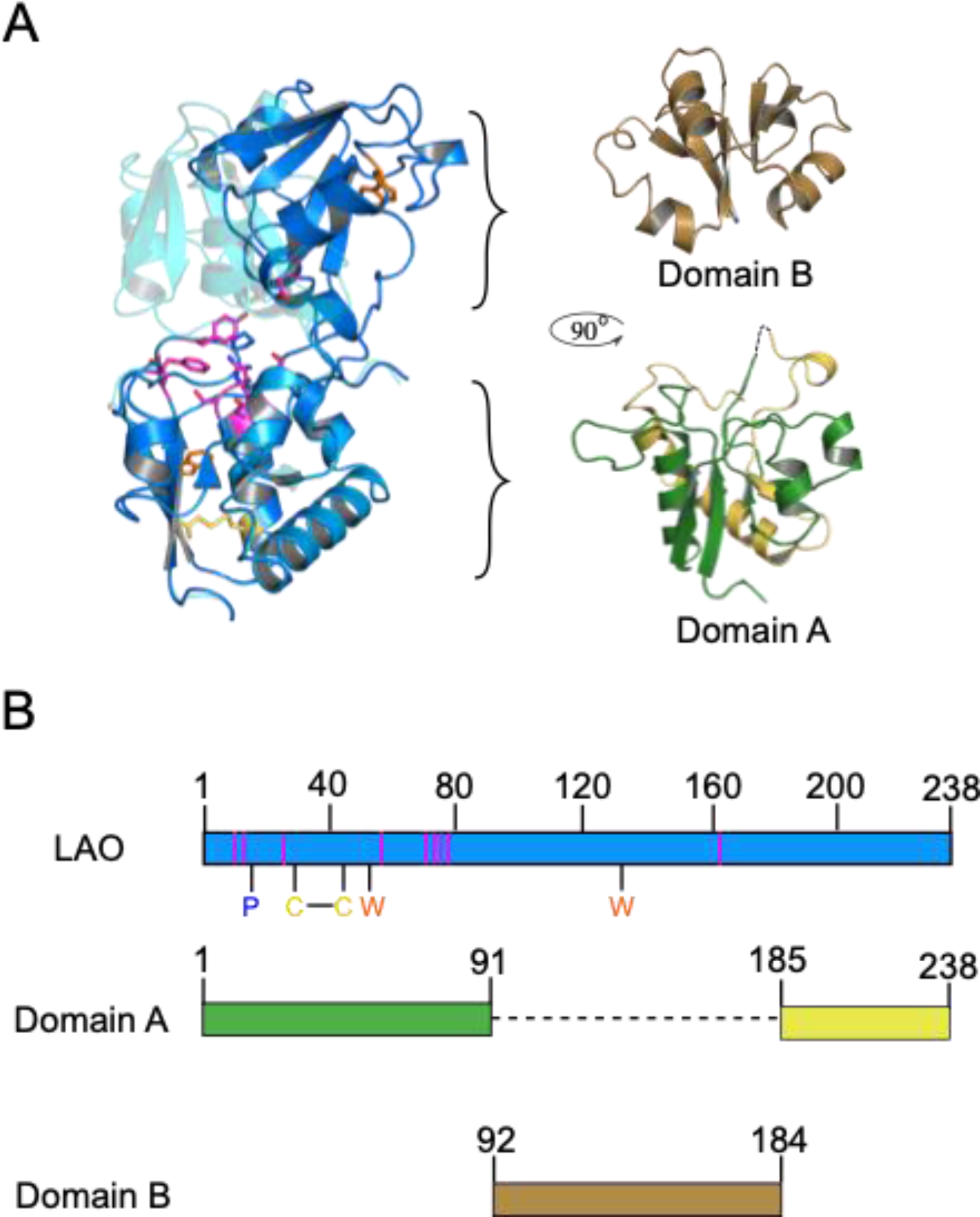
Structure of LAO and schematic representation of the individual domains A and B. A) Structural alignment of LAO in open (blue, PDB ID = 2LAO) and closed (cyan; PDB ID = 6MLE) states. Residues 1-80 and 190-238 were aligned for the superposition. W47 and W130 (orange), P16 (blue), the nine binding site residues (magenta) and the disulfide bond formed by the residues C38 and C45 (yellow) are depicted as sticks. The continuous (brown) and discontinuous (green and yellow for the N-terminal and C-terminal elements) domains of LAO are shown independently. B) Schematic representation of the sequences of LAO and the independent domains. The domain A was produced by expressing the residues 1-91 and 185-238 without an additional linker in between; the domain B expressed individually comprises residues 92-184. Lines in the LAO scheme show the position of the aforementioned residues.

## Results

Far-UV Circular Dichroism (CD) was used to monitor changes in the secondary structure of LAO and its two independent domains. The CD spectrum of LAO in native conditions (pH 9.0) is typical of αβ proteins, with minima at 210 and 222 nm, signature of α structure, moderated by the 218 nm signal, characteristic of β-sheet. (Figure 2A). A single chain version of the discontinuous domain A, thereafter, referred to as dA, was constructed by joining the hinge residues 91 and 185 (Figure 1B) which corresponds to 61% of the sequence of LAO. In addition, the independent domain B, hereafter named dB, was constructed by expressing the continuous residues 92 to 184 (Figure 1B). Although dA is assembled from two discontinuous segments, it shows the characteristic features observed in the CD spectrum of the complete folded protein, with a molar ellipticity proportional to the size of the domain (Figure 2B). By contrast, the shape and signal of the CD spectra of dB are both consistent with an unfolded state at pH 9.0 that becomes structured in more acidic conditions (Figure 2C). dB was therefore studied at pH 5.0. The sum of the CD spectra of dA plus dB resembles the spectra of LAO even when recorded at different pH (9.0 and 5.0, respectively). From these observations it can be noted that the secondary structure of each independent domain seems to be like that of its corresponding segments in the complete protein (Figure 2A).

**Figure 2.**
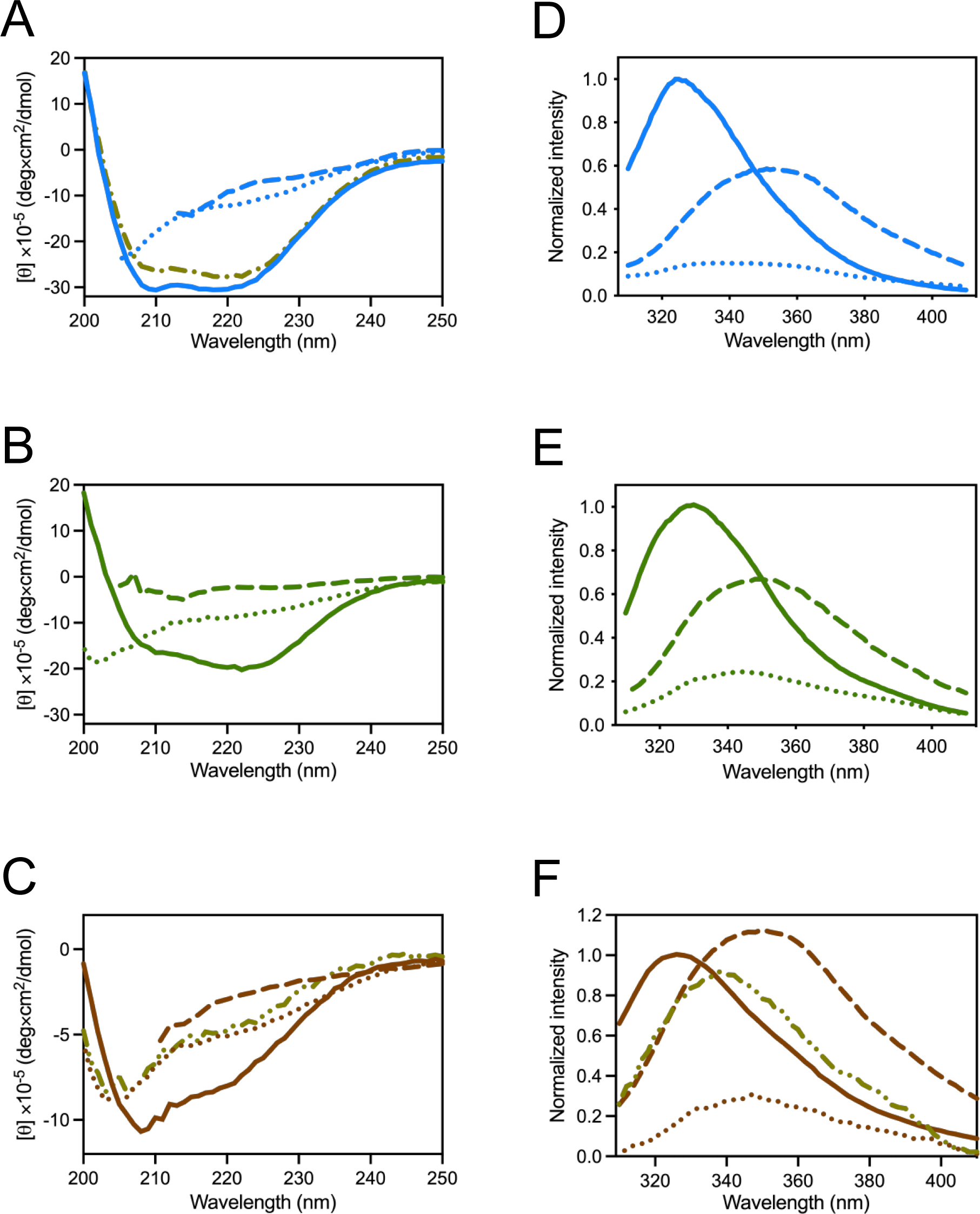
Spectroscopic characterization of LAO and its individual domains. Circular dichroism and fluorescence emission spectra of LAO (A and D, respectively, blue lines) and dA (B and E, respectively, green lines) at pH 9.0, and dB (C and F, respectively, brown lines) at pH 5.0. Spectra were recorded at 25 °C (solid lines), 80 °C (dashed lines) and 25 °C in the presence of 6 M urea (dotted lines). Dash-dotted olive lines show the sum of the spectra of folded dA (pH 9.0) and dB (pH 5.0) at 25 °C in panel A and the spectra of dB at 25 °C, pH 9.0 in panels C and F.

Intrinsic fluorescence spectroscopy (IF) was used to follow the tertiary structure. LAO contains two tryptophan residues, W47 and W130, partially buried from the solvent in dA and dB, respectively (Figure 1A and B). The IF spectra of LAO, dA and dB in native conditions show intensity maxima around 325 nm (Figure 2D, 2E and 2F). In LAO and dA this peak shifts to a higher wavelength upon thermal or urea-induced unfolding, reflecting the complete exposure of tryptophan residues to the solvent (Figure 2D, and 2E). Further use of fluorescence spectroscopy to follow the unfolding of dB was hampered due to the small intensity changes observed in chemical unfolding and the severe quenching observed in temperature-induced unfolding (Figure 2F).

The thermal unfolding of LAO and dA was explored at pH 9.0 due to irreversibility of the process at pH values lower than 8.0. The unfolding of dB was studied at pH 5.0 since, as mentioned above, this domain is unstable at pH 9.0. The temperature-induced unfolding of LAO showed monophasic and coincident transitions when followed by CD and IF (Figure 3A and Figure S1A). Likewise, DSC endotherms were highly symmetric and showed a single transition (Figure 3B and Figure S1B). Endotherm recovery in a second DSC scan varies between 86% and 96%, evidencing the high reversibility of LAO unfolding (Table 1). A two-state equilibrium model properly fitted spectroscopic and calorimetric data (Figure 3A and B and Figure S1A and B). Further evidence for the two-state character of LAO unfolding comes from the similarity in the Tm values obtained by spectroscopic and calorimetric data, as well as the calorimetric ratio (ΔHcal/ΔHvańt Hoff = 1.0) (Table 1).

**Figure 3.**
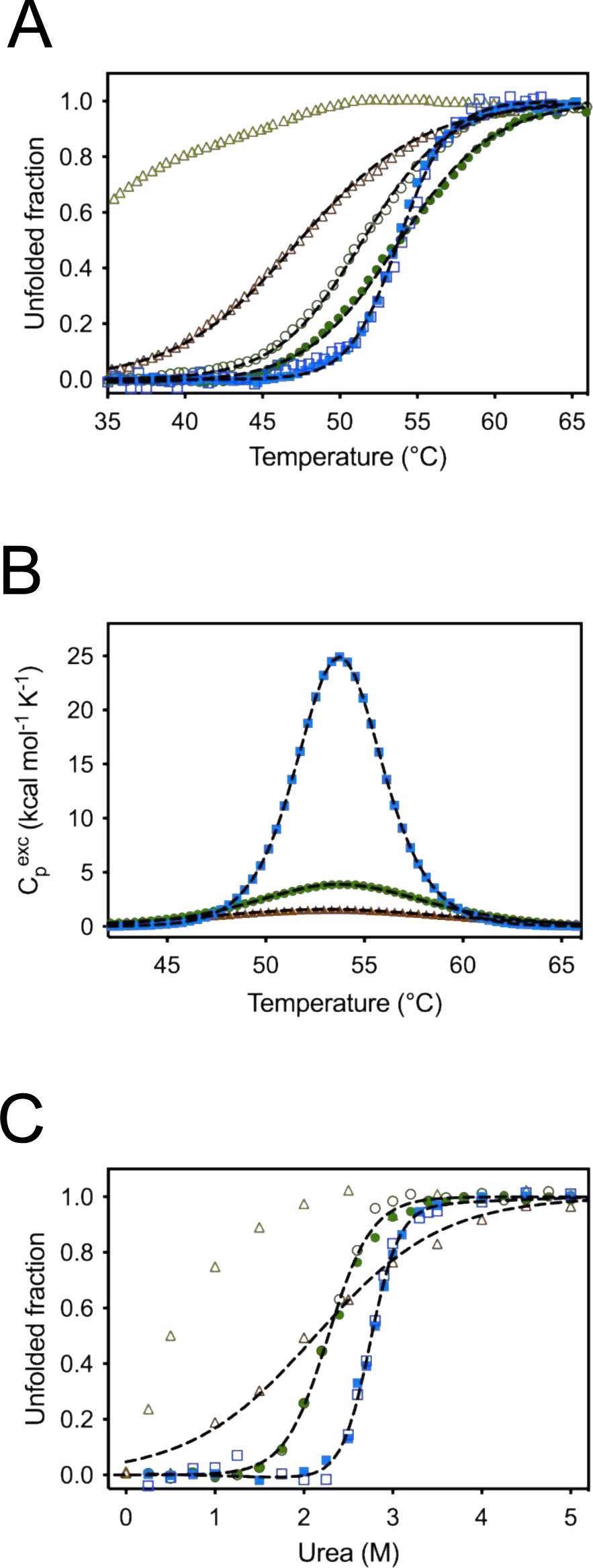
Thermal and urea-induced unfolding of LAO and its isolated domains. A) Unfolded fraction curves measured by circular dichroism (open symbols) or fluorescence intensity (closed symbols). The fraction of unfolded protein of LAO (blue) and dA (green) at pH 9.0, as well as dB at pH 5.0 and pH 9.0 (brown and light green, respectively) were fitted to a two-state model (black lines). B) Differential scanning calorimetry traces of LAO and dA at pH 9.0 (blue and green, respectively) and dB at pH 5.0 (brown); the baseline-subtracted curves for LAO and dA were fit to a two-state model while the curve for dB was fitted to a non-two-state model where the Van’t Hoff and calorimetric ΔH are fitted independently (black lines). C) Urea-induced unfolding curves measured by circular dichroism (open symbols) and fluorescence intensity (closed symbols). The fraction of unfolded protein of LAO and dA at pH 9.0, as well as dB at pH 5.0 and pH 9.0 were fitted to a two-state model (black lines). The color code is as described for panel A.

**Table 1.** Thermodynamic parameters for temperature and urea-induced unfolding.

| pH | Temperature-induced unfolding |  |  |  |  |  | Urea-induced unfolding |  |  |
| --- | --- | --- | --- | --- | --- | --- | --- | --- | --- |
|  | DSC |  |  | Spectroscopic |  | <sup>a</sup>(ΔH<sub>cal</sub>/ΔH<sub>vh</sub>) | Spectroscopic |  |  |
|  | T<sub>m</sub><br>(°C) | ΔH<sub>cal</sub><br>(kcal mol<sup>-1</sup>) | Reversibility<br>(%) | T<sub>m</sub><br>(°C) | ΔH<br>(kcal mol<sup>-1</sup>) |  | ΔG<br>(kcal mol<sup>-1</sup>) | *m*<br>(kcal mol<sup>-1</sup> M<sup>-1</sup>) | C<sub>m</sub><br>(M) |
| LAO |  |  |  |  |  |  |  |  |  |
| 9.5 | 52.8 ± 0.1 | 144.2 ± 0.4 | 96 | 52.6 ± 0.1 | 136.7± 4.1 | 1.05 ± 0.03 |  |  |  |
| 9.0 | 54.3 ± 0.1 | 145.8 ± 0.5 | 96 | 54 ± 0.1 | 140.7 ± 4.0 | 1.04 ± 0.03 | 9.8 ± 0.8 | 3.4 ± 0.3 | 2.8 ± 0.1 |
| 8.5 | 55.5 ± 0.1 | 151.7 ± 0.3 | 86 | 55.6 ± 0.1 | 143.7 ± 6.2 | 1.03 ± 0.04 |  |  |  |
| 8 | 57.1 ± 0.2 | 153.0 ± 0.6 | 86 | 57.0 ± 0.1 | 147.4 ± 5.8 | 1.04 ± 0.04 | 12.1 ± 2.3 | 3.8 ± 0.7 | 3.2 ± 0.3 |
| dA |  |  |  |  |  |  |  |  |  |
| 9.0 | 53.2 ± 0.1 | 71.4 ± 0.2 | 79 | CD<br>51.4 ± 0.3<br>IF<br>53.8 ± 0.2 | CD<br>75.4 ± 6.6<br>IF<br>71 ± 2.4 | 0.95 ± 0.04 | 5.5 ± 0.2 | 2.4 ± 0.1 | 2.3 ± 0.1 |
| dB |  |  |  |  |  |  |  |  |  |
| 5.0 | 53.6 ± 0.1 | <sup>b</sup>27.2 ± 1 | 96 | 47.4 ± 0.1 | 51.2 ± 2 | 0.52 ± 0.04 | 1.8 ± 0.2 | 0.8 ± 0.1 | 2.2 ± 0.3 |

<sup>a</sup>Ratio calculated from the calorimetric enthalpy ( $\Delta H_{cal}$ ) obtained from DSC and the van't Hoff enthalpy ( $\Delta H_{vh}$ ) from spectroscopic data.
<sup>b</sup>For this condition the “non-two state model”, provided in the Origin software, where $\Delta H_{cal}$ and $\Delta H_{vh}$ are calculated independently, was used to fit DSC data. $\Delta H_{vh}$ obtained from calorimetric data for dB was 52.5 kcal mol<sup>-1</sup>.

The thermal unfolding of dA followed by CD, IF and DSC showed single transitions that were properly fitted to a two-state model (Figure 3 A, 3B and Table 1). A difference of ∼ 2 °C was observed between the Tm values calculated from CD and IF data (Table 1), suggesting an asynchronous loss of secondary and tertiary structure. dA contains ∼60 % of the complete protein residues; in accordance, the unfolding ΔH of this domain corresponds to ∼50 % of that for LAO. The temperature-induced unfolding of dB showed a broad transition when followed by CD (ΔHvańt Hoffl= 51.2 kcal mol^−1^), with a Tm that clearly differs by ∼ 6 °C from the one obtained by DSC. In this case, proper fitting of the endotherm required the independent estimation of ΔHcal and ΔHvańt Hoff, indicating the presence of a folding intermediate (Table 1, see materials and methods). The ΔHcal value observed for dB was very low and not proportional to its size. These data show that dB deviates substantially from its native-like conformation in the absence of interdomain interactions.

CD and IF addressed the urea-induced unfolding of LAO and its isolated domains to further explore its behavior at equilibrium in the presence of chemical denaturants and determine its stability at room temperature. Transitions followed by both techniques were all monophasic and fitted properly to a two-state model (Figure 3C). The ΔG of LAO and dA are proportional to their size, while dB is marginally stable (Table 1). Overall, the ΔH and ΔG values of the isolated domains do not add up to the thermodynamic properties of the complete protein.

### Unfolding and refolding kinetics of LAO

In order to gain insight into the folding mechanism of LAO, stopped-flow experiments monitored by CD and IF were performed. All unfolding kinetic curves showed a single phase (Figure S2), with rate constants linearly dependent on urea concentration in a chevron plot (Figure 4A). Refolding kinetics were also monophasic at urea concentrations higher than 1.5 M (Figure S3), whereas biphasic kinetics were observed below 1.5 M urea (Figure S4). Consequently, two branches are present in the refolding side of the chevron plot (Figure 4A). The dependence of both refolding rate constants on urea concentration shows a curvature. This behavior, together with the presence of slow and fast refolding rates, have been ascribed to the presence of kinetic intermediates and/or to the *cis-trans* isomerization of proline or non-proline residues [45–53].

**Figure 4.**
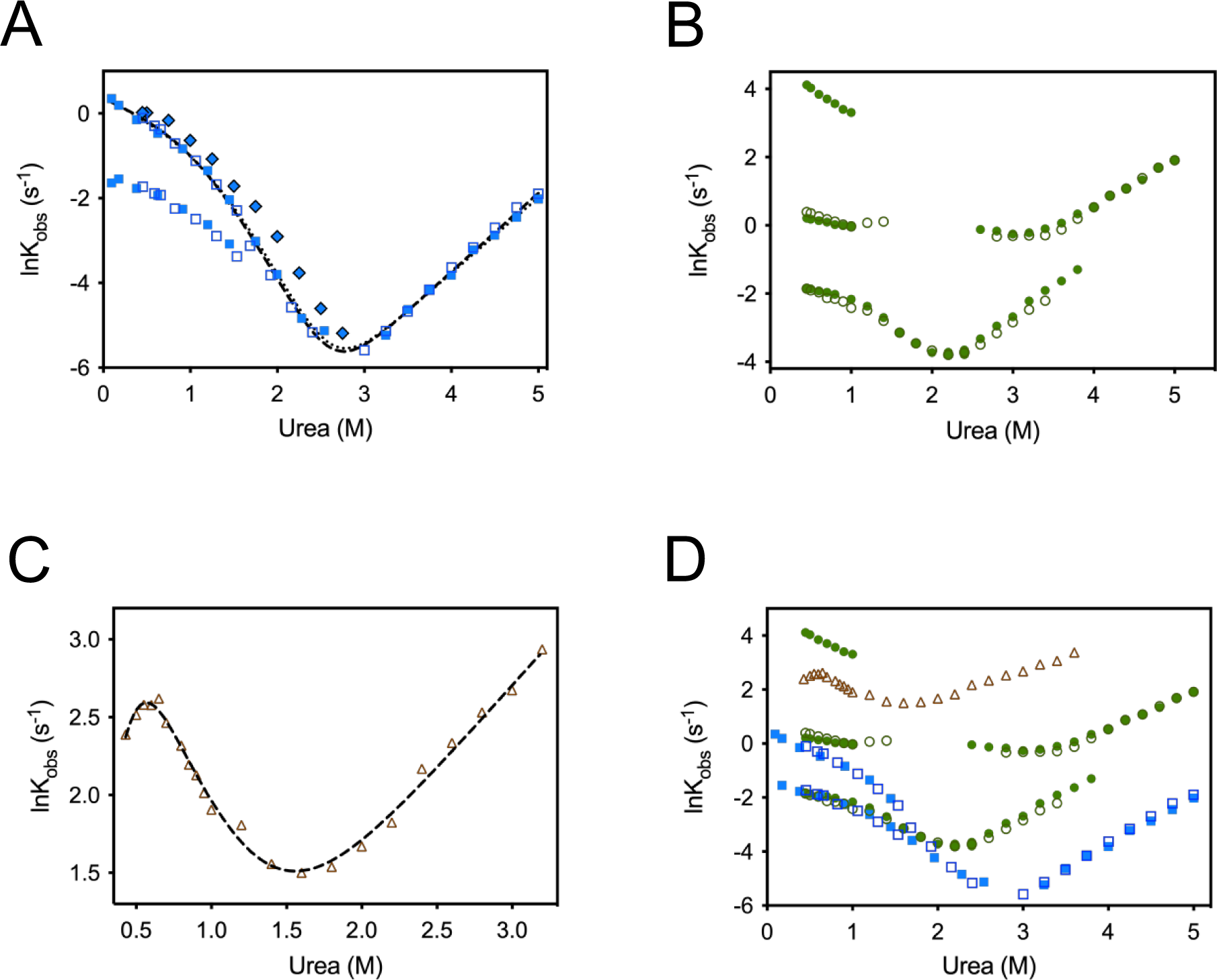
Chevron plots of LAO, dA and dB. A) Chevron plot of LAO obtained from the unfolding and refolding kinetics measured by circular dichroism (open symbols) and fluorescence intensity (closed symbols). The fast refolding and unfolding branches were fit to three-state models with on-pathway (dashed line) or off-pathway (dotted line) intermediates. The closed diamonds show the refolding branch obtained in the presence of 5 uM cyclophilin. B) Chevron plots of dA obtained from the unfolding and refolding kinetics measured by circular dichroism (open symbols) and fluorescence intensity (closed symbols). C) Chevron plot of dB obtained from the unfolding and refolding kinetics measured by circular dichroism. Data were fitted to a three-state model with a non-productive intermediate (Methods, scheme 2). D) Comparison of the Chevron plots of LAO, dA and dB according to the color scheme shown in panels A to C.

The contribution of proline-peptide bonds isomerization to refolding kinetics can be assessed in “interrupted unfolding” experiments, also known as “double-jump” assays [54]. In the first jump, the protein is unfolded at a high concentration of denaturant and then refolded in the second jump by dilution in the working buffer. Refolding amplitudes increase with the first jump length due to the appearance of the unfolding state; nevertheless, since backbone isomerization is a slow process (relaxation times ∼ 10s or higher at 25 °C [55], its contribution should be small at short unfolding times. LAO’s interrupted unfolding experiments did show such behavior; indeed, a single exponential describes closely the refolding kinetics at short unfolding times (≲ 3 s) (Figures S5A and S6). From this evidence the slow phase observed in refolding kinetics might be due to proline isomerization. This was further explored by carrying out refolding experiments in the presence of the *cis-trans* prolyl-isomerase cyclophilin A (CyP). Monophasic kinetics were detected when CyP was added (Figure S5B) and the refolding branch of the chevron plot obtained in the presence or absence of CyP are very similar (compare rhombi and squares in Figure 4A). The curvature is observed in both refolding arms, indicating that an intermediate is populated with or without CyP.

To further characterize the folding mechanism of LAO, the fast refolding and unfolding branches of the chevron plots were fitted to both on-pathway (sequential mechanism) and off-pathway (non-productive mechanism) models (See schemes 1 and 2, respectively in methods). Experimental data are adequately described by both mechanisms (Figure 4A, Table 2); accordingly, interrupted refolding experiments were carried out to obtain evidence of the time evolution of different folding species [56]. This type of experiment consists of two steps: 1) unfolded LAO is mixed with buffer to allow refolding; 2) the protein is mixed with a concentrated urea solution to promote unfolding. The time lapse between the first and second steps is varied to allow refolding to reach different extents, then the unfolding kinetics of the second step are recorded. The number of rate constants required to fit the unfolding curves is related to the number of transitions between states, whereas the amplitudes are proportional to their concentration. All kinetic traces were biphasic (Figure S7), indicating that at least two states are populated during refolding. The rate constant estimated for the slow phase is very similar to the single constant obtained from “normal” unfolding kinetics (Figure S2), suggesting that this phase corresponds to the unfolding of the native state. The time evolution of the amplitude associated with the faster rate shows the characteristic behavior of an intermediate (Figure S8). Amplitudes can be adequately fitted to both the on-pathway and triangular mechanisms (Schemes 1 and 3, Figure S8 A and C), while the off-pathway (Scheme 2) can be clearly ruled out (Figure S8B). Although these two single-intermediate mechanisms can properly fit the observed data, they are not truly consistent with all the observations described above, especially with the presence of a rate limiting step related to proline isomerization. The values for the rate constant *k_f1_* obtained from data fitting to schemes 1 and 3 (0.17 and 0.12 s^−1^ respectively, Table S1) are very similar to the one obtained for the slow branch in the chevron plot at the same condition (0.18 s^−1^ at 0.45 M urea). This suggests that the interconversion between I and N might be limited by proline isomerization.

**Table 2.**
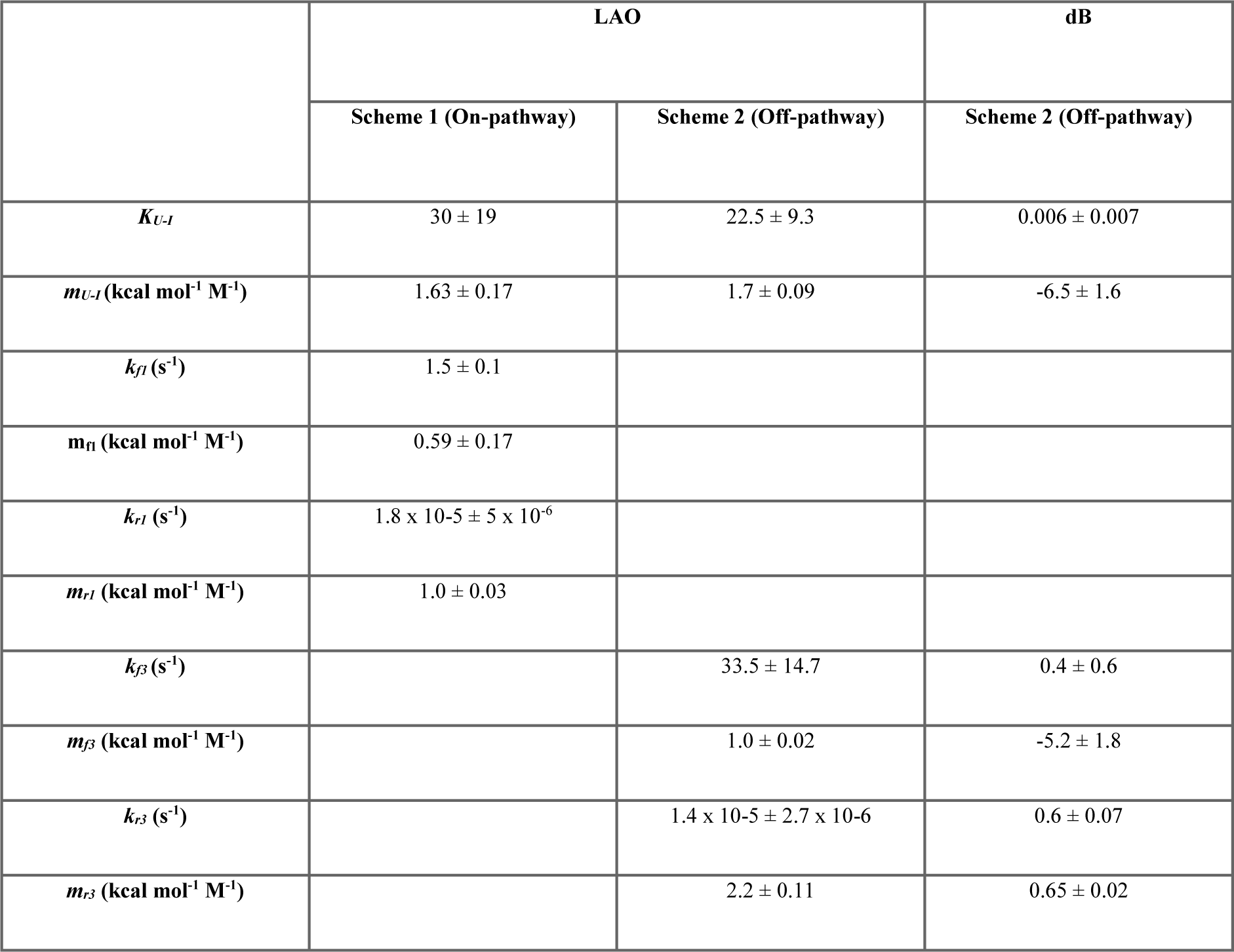
Kinetic parameters from Chevron plots

The simplest scheme compatible with all the experimental observations is expected to include at least two pathways, each of them with one intermediate. Scheme 4 depicts such a model, where I_C_ represents the ensemble of intermediates with all the proline residues matching the configuration observed in the native state. In contrast I_T_ represents all the conformers having at least one proline residue in a non-native configuration. This scheme properly describes the time evolution of the amplitudes in the interrupted refolding experiments, as observed in Figure S8. However, the analysis of the rate constants (Table S1) predicts a slow interconversion rate between U and I_C_. When this last step is neglected, the system reduces to a branched mechanism with a dead-end intermediate (Scheme 5), which also describes experimental data. In this case, the role of I_T_ is difficult to explain since the experimental evidence points to a proline *cis*-*trans* isomerization as the limiting step. However, the interconversion between I_T_ and I_C_ can be envisaged as a simplification of a more complex process that cannot be resolved during kinetics, as depicted in the scheme 6. As expected for the increase in the number of fitting parameters, data can also be described by this complex mechanism, however, some estimates for the rate constants are ill-defined, due to their large uncertainties which are a consequence of the system’s overdetermination (Table S1).

### Unfolding and refolding kinetics of the isolated domains A and B

Unfolding assays of dA showed single exponential kinetics when measured at urea concentrations higher than 3.75 M (Figure S9). By contrast, a second exponential was present at lower denaturant concentrations (Figure S10), resulting in two unfolding branches in the chevron plot (Figure 4B). Both rate constants are faster than those observed for the unfolding of LAO (Figure 4D). Refolding kinetics measured below 2.0 M urea, followed by CD, were properly fitted by two exponentials, whereas those monitored by IF required three exponentials (Figure S11). This complex behavior results in three refolding branches in the chevron plot of dA (Figure 4B). Two of these branches overlap with those obtained for LAO (Figure 4D), including the slowest one, related to proline isomerization in the complete protein. In addition, a burst phase associated with a decrease in fluorescence intensity was detected in the refolding kinetics of dA (compare the final IF signal in Figure S9B with the initial signal in Figure S11B). A similar behavior has been previously attributed to the formation of a compact intermediate state [57,58]. The fastest IF constant observed after the burst phase, corresponding to the increase in intensity that characterizes unfolding kinetics (Compare Figure S10B and S11B), should be related to the relaxation of this compact state. Due to the complexity observed in the kinetics of dA, we were unable to propose a particular folding mechanism for this domain; nevertheless, judged from the kinetic data, it might involve three or more distinct folding intermediates.

In contrast to dA and LAO, dB unfolding, and refolding can be properly described by single exponentials (Figure S12). A downward curvature in the dB chevron plot is considered characteristic of an off-pathway intermediate. (Figure 4C). The rate constants for the unfolding and refolding of dB were faster than those observed for LAO and dA (Figure 4D) and no burst or refolding bifurcation were observed.

### Structural and functional characterization of dA

Though the spectroscopic signals gave evidence of a similar native fold for dA, when isolated or as a domain of LAO, further details of the structural impact of removing the interdomain connections were obtained by solving the structure of this domain (Figure 5 and Table S2). The X-ray crystallographic structure of the isolated dA superimpose with a very low RMSD (0.3 Å-124 Cɑ) with the domain A of LAO (Figure 5A). The extensive interactions between secondary structure elements from both the N- and C-termini of LAO are preserved in the structure of dA; the only minor differences were observed in the region where both termini were fused (Figure 5B). The side chain conformation of the residues at the putative binding site in dA resemble the open state of LAO (Figure 5C). By contrast, when dA is compared to the closed state, differences are found in the conformation of the binding site residues Y14 and S72 (Figure 5D); therefore, dA is a good model for the open state of the complete protein.

**Figure 5.**
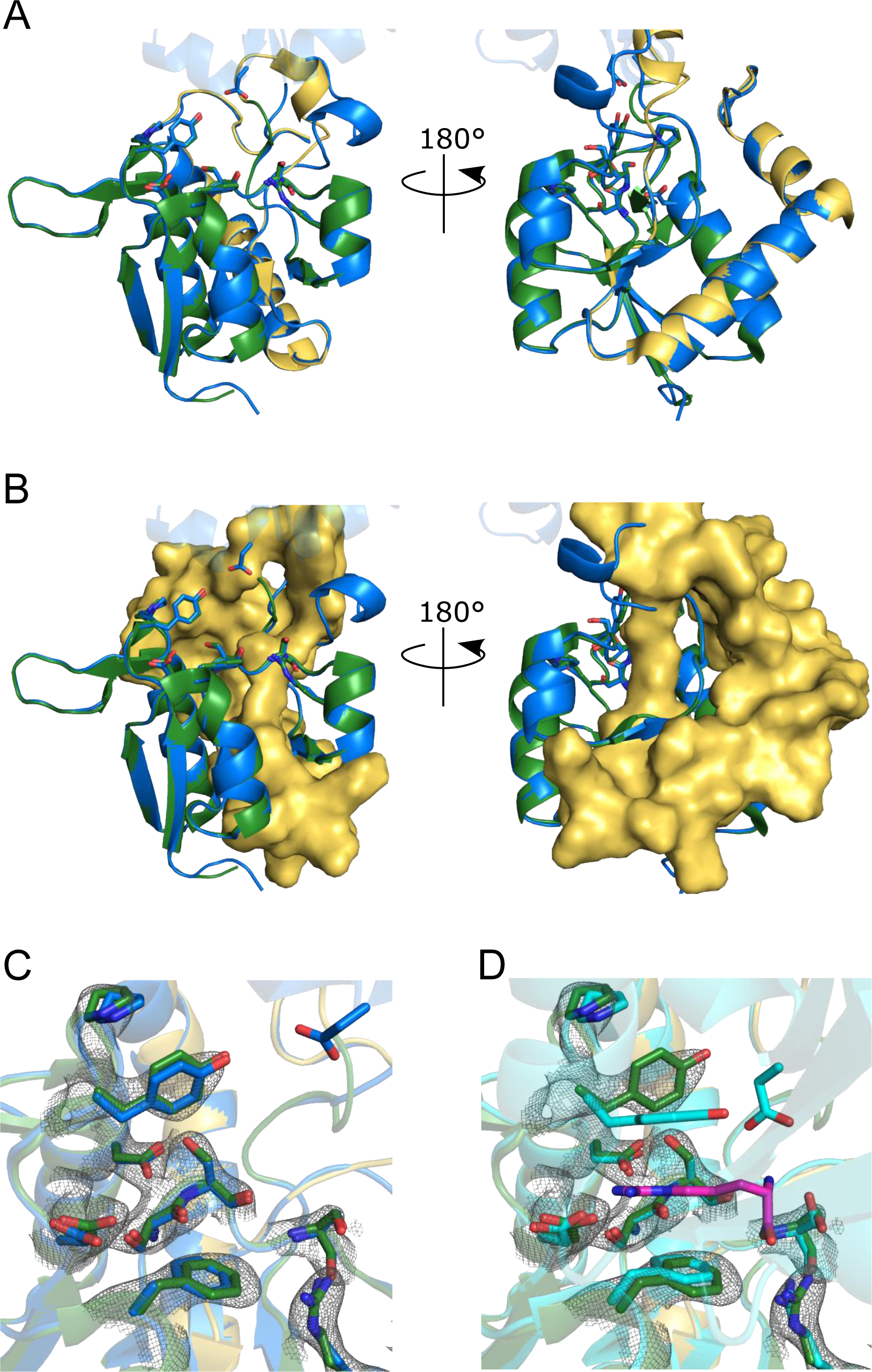
Crystallographic structure of dA in absence of ligand. A) Structural alignment of dA (green and yellow for the N-terminal and C-terminal halves, respectively; PDB ID = 6XKS) and the corresponding domain of LAO in the open state (blue) (RMSD = 0.3 Å-124 Cɑ-). The binding site residues are shown as sticks. B) The C-terminal half of dA is shown as a surface to show that the high complementarity between both halves of the protein is maintained in the individual domain. C) and D) Close-up of the protein binding site in the structural alignment of dA and the open (C, blue) and closed (D, cyan) states of LAO. The 2fo-fc electron density map is shown for the binding site residues of dA as a gray mesh while the arginine ligand is presented in D as magenta sticks. The conformation adopted by the binding site residues of dA corresponds to that observed in the open state of LAO.

The functional abilities of dA were then addressed by ligand binding experiments followed by titration calorimetry experiments. In LAO, arginine binding is an enthalpy-driven process (ΔHLAO= −11.3 kcal mol^−1^) with a slightly favorable entropy change (-TΔSLAO = −1.0 kcal mol^−1^), that exhibits nanomolar affinity (KD= 1.0 nM) [41]. ITC experiments reveal that arginine binding to dA is less exothermic when compared to LAO (ΔHdA= −7.7 kcal mol^−1^) and becomes entropically unfavorable (-TΔSdA = +2.5 kcal mol^−1^) (Figure 6). Consequently, the affinity of dA for arginine is severely reduced (KD= 131 µM).

**Figure 6.**
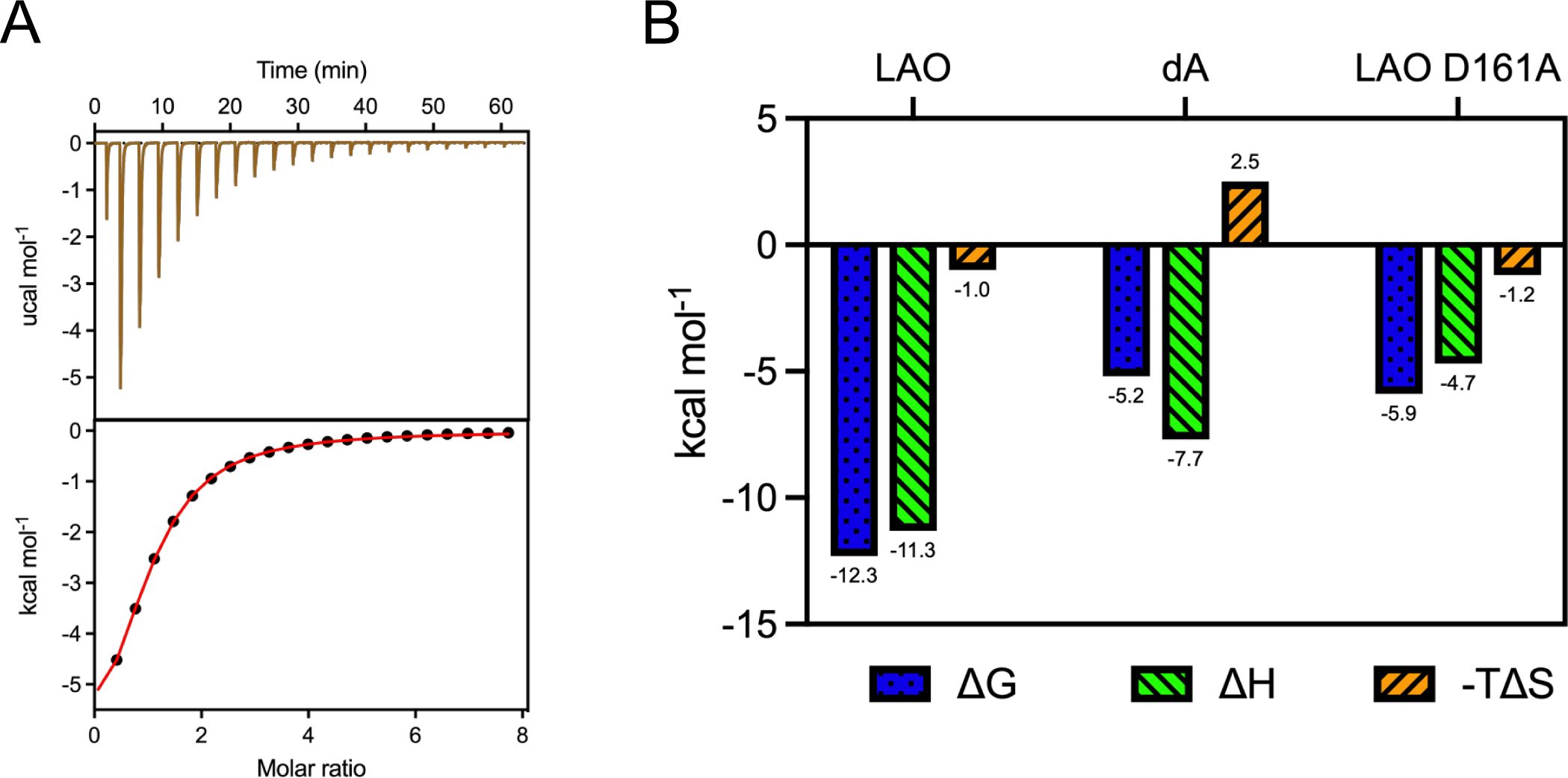
Binding thermodynamics of LAO, dA and the mutant D161A. A) ITC experiment of dA titrated with L-arginine; the upper panel displays the heat released following each injection after baseline subtraction while the lower panel shows the integration heats fit to a 1:1 model. B) Comparison of thermodynamic signatures for arginine binding to LAO, dA, and the LAO D161A mutant, where the only residue of the binding site located in dB has been removed.

The structural and energetic role of ligand binding to LAO has been previously studied by alanine scanning mutagenesis of the nine binding site residues [43]. Most of them are located in domain A and only D161 is placed in domain B, therefore, the number of binding site residues of the mutant D161A and dA are equivalent. The arginine binding event in D161A is moderately exothermic (ΔHD161A= −4.7 kcal mol^−1^) and entropically favorable (-TΔSD161A= −1.2 kcal mol^−1^), resulting in a substantial drop in affinity (KD = 42 µM) [43]. As expected from the similar number of binding residues present in dA and D161A, both proteins show similar binding energy (ΔΔGdA-LAOD161A= 0.7 kcal mol^−1^). However, ligand binding is less exothermic (ΔΔHdA-LAOD161A=-3.0 kcal mol^−1^) and more entropic (Δ(TΔS) dA-LAOD161A= 3.7 kcal mol^−1^) in dA when compared to D161A. These differences in the thermodynamic signatures suggest that solvent release upon ligand binding is less extensive in dA than in LAO and D161A due to the lack of a conformational change in dA [41,43].

## Discussion

The characterization of multi domain proteins is required to advance in the understanding of protein folding and function in complex systems. Here, we addressed the folding and ligand binding of the bilobal protein LAO in comparison to the behavior of its two isolated domains. Both equilibrium and kinetic experiments show that LAO unfolding/refolding is a highly reversible process which is then suitable for an in-depth physicochemical characterization. As judged from spectroscopic and calorimetric data, both the thermal- and urea-induced unfolding of LAO are two-state processes. The absence of stable intermediates indicates a highly cooperative unfolding of both domains, as observed in other PBPs such as MBP and ABP [31,34]. On the one hand, given the discontinuity in the amino acid sequence of the domain A, we anticipated profound alterations in the stability and structure of dA. Nevertheless, this domain adopts the same structure when isolated or when integrated into LAO and exhibits a stability that corresponds to that expected for its size. On the other hand, given the continuous nature of the domain B sequence, we assumed that there would be no significant structural and energetic disturbances in the isolated domain. Notwithstanding, dB was barely stable when expressed independently. These findings reveal that, opposite to our initial hypothesis, the stability of the continuous domain is highly dependent on the presence of the discontinuous domain which does not require the presence of its counterpart. Overall, the sum of the stabilities of dA and dB is lower than the stability of LAO, revealing that the structural domains of LAO are thermodynamically coupled. Previous studies on the unfolding of multidomain proteins have led to the hypothesis that proteins with large domain interfaces are stabilized by their neighbors and fold cooperatively. Conversely, multidomain systems with short flexible linkers and small interfaces, tend to behave as independent units [25]. The results described above show that LAO is a clear exception to this trend, because in the absence of ligands, the interaction between domains is limited to the small hinge region. These observations suggest that other factors such as the open/close equilibrium and topological constraints are involved.

In contrast to the simplicity observed in equilibrium studies, kinetic experiments unveiled a complex folding mechanism. The results obtained from the detailed kinetic characterization of LAO indicate a folding process best described by a mechanism involving at least two intermediates in parallel branches. As judged from the effect of cyclophilin A, and double jump experiments, one of them is most likely related to *cis-trans* proline isomerization of a peptide bond. It should be noted that of the seven proline residues present in LAO, four of them are located in dB and the other three in dA, including P16. This proline is functionally relevant and conserved in PBPs [59], and is the only residue that adopts a *cis*-peptide bond in the native state of both LAO and dA. Since one of the three refolding branches observed in the chevron plot of dA is equivalent to the refolding arm of LAO that disappears in the presence of cyclophilin, it is reasonable to ascribe the *cis-trans* proline isomerization process observed in LAO to P16.

The folding behavior of LAO and their isolated domains differ considerably. The continuous domain dB follows a relatively simple folding mechanism, with one off-pathway intermediate not observed in LAO. Conversely, dA shows a very complex mechanism that seems to include four or more intermediates, one of them resulting in a burst phase. Because the overall structure of dA and domain A superimpose, this intricate behavior must be a consequence of the remodeled connectivity rather than rearrangements of native interactions, as has been described in other systems [60–62]. Although small systems might be expected to exhibit a simpler folding behavior than larger ones, the folding of dA is more complex than the observed for the complete protein. A complex folding mechanism has been also reported for the 14 kDa protein CheY, that shows similar topology than dA and dB. The characterization of both CheY and dA show evidence of parallel folding pathways associated with proline isomerization and misfolded intermediates [63]. Indeed, it has been proposed that periplasmic binding proteins arose from gene duplication of a CheY-like protein [38]. This evolutionary relation between CheY and PBPs along with the similarities in the folding behavior of dA and CheY confirm the finding that, even in the absence of other intramolecular interactions, the Rossmann fold is topologically frustrated [63]. The complex folding behavior of CheY has been related to the presence of non-native interactions during the folding process. A reasonable hypothesis for the observed simplicity in the folding mechanism of LAO is that the presence of the continuous domain, that also folds and unfolds via a simpler mechanism, introduces constraints in the folding process of the complete protein that avoids non-native interactions. This is in line with the suggestion that multidomain proteins have evolved mechanisms that avoid misfolding events due to interdomain interactions [25]. Since PBPs are present in the three domains of life [35,36], the gene duplication event that produced them should be ancient. Therefore, the functional unit of LAO has a long evolutionary history and there seems to be no evolutionary pressure to maintain the stability and cooperativity of the isolated lobes, instead, both domains may have evolved as a single component whose folding and function are also linked.

The first structural and functional characterization of the individual lobes of a PBP was carried out on HisJ [44]. LAO and HisJ share 70 % sequence identity, are codified by adjacent genes, interact with the same membrane transporter (HQM2), and bind similar ligands but with an opposite order of affinities [32,64]. The isolated domains of HisJ were successfully expressed and their NMR characterization showed that, at pH 7.0, both fold into equivalent structures when isolated or covalently linked. This does not correlate with the behavior observed in LAO, where dB is not properly folded at similar conditions (pH ≥ 7.0) and becomes marginally stable in more acidic conditions. Regarding interaction with their preferred ligand, both histidine binding to HisJ and arginine binding to LAO exhibit nanomolar affinity (114 nM and 1 nM, respectively) and enthalpy-driven ligand binding (−4.6 and −11.3 kcal mol^−1^, respectively) [41,44]. Interestingly, the independent discontinuous domains of both proteins show very similar binding affinities (80 and 154 µM for dA-LAO and d1-HisJ, respectively), while the continuous domain shows no evidence of binding. Despite these similarities, the thermodynamic signatures of the independent discontinuous domains are very different: whereas in dA-LAO ligand binding is still very exothermic (−7.7 kcal mol^−1^), the enthalpy change associated to ligand binding in d1-HisJ was below calorimetric detection, indicative of an entropy-driven process. The structures of both LAO and HisJ in the presence of ligands show that eight out of the nine direct protein-ligand interactions come from their discontinuous domains. Therefore, it is not surprising that dA-LAO and d1-HisJ can interact with their ligands whereas ligand binding to dB-LAO and d2-HisJ was not detected. Due to the detection limits of ITC and NMR, ligand binding to the continuous lobes is at best in the low micromolar range, corresponding to a maximum binding energy on the order of ∼3 kcal mol^−1^. If ligand binding to PBPs were additive, the ΔG calculated from the sum of the values observed for the isolated lobes would be ∼8 kcal mol^−1^. However, the observed ΔG values are ∼12 kcal mol^−1^, thus the binding energy cannot be explained just as the sum of the two independent domains and exceeds simple addition by ∼4 kcal mol^−1^, indicating a strong coupling of both domains for the energetics of ligand binding.

In conclusion, we have shown further evidence on how the bilobal nature of PBPs contribute to their conformational stability, folding cooperativity and binding affinity, in a manner exceeding the simple sum of the contribution of each domain taken independently. Since the bilobal nature of SBPs is the outcome of an ancient domain duplication event [38], it seems that both domains have coevolved to fold into a stable and functional single entity.

## Methods

### Protein expression and purification

The gene of the independent domain A was generated by PCR from the gene encoding for LAO, cloned into pET12b. LAO and dA contain one N-terminal signal sequence required for transport to the periplasm, here it is cleaved upon arrival to produce the mature protein. This allows the use of the osmotic shock protocol for the expression and purification of these proteins as described in Vergara et al. [43]. Briefly, pET12b vectors containing the corresponding genes were transformed into *Escherichia coli* cells of the strain BL21-A1. Transformed cells were grown in LB agar plus ampicillin and single colonies were used to inoculate 10 mL of LB media supplemented with 100 µg/mL of ampicillin that were then incubated overnight at 37 °C. Starter cultures were transferred to 1L of TB media plus 100 µg/mL of ampicillin and were incubated at 37 °C until reaching and OD = 0.8-1.0. Protein expression was induced with 0.25% of L-arabinose for 4 h, at 37 °C. Cells were harvested by centrifugation at 5129 g for 10 min, and pellets were resuspended in a solution of potassium acetate 10 mM, at pH 5.1, with 20% sucrose and 1 mM EDTA. Cells were harvested for a second time and pellets were gently resuspended in a solution of potassium acetate 10 mM, at pH 5.1, to promote the release of the protein by osmotic shock. After centrifuging at 15253 g for 20 min, the supernatant was collected and concentrated. The protein was unfolded to remove bound ligands by adding a solution of Tris 5 mM, at pH 8.5, with guanidinium chloride (GdnHCl) 2M, and centrifuging in 10 K (3 K for dA) ultra-centrifugal filters Amicon® (Merck KGaA, Darmstadt, Germany). This process was repeated five times to dialyze the ligand. GdnHCl was removed to promote refolding by adding buffers without denaturant and centrifuging for at least five times. Finally, the protein sample was loaded into a HiTrapTm Q HP anion exchange column (GE Healthcare Bio-Sciences, Pittsburgh, PA, USA) and eluted with ∼ 50 mM NaCl, using a 0-500 mM linear gradient.

The individual domain B was purified following the His-tag purification protocol to obtain the required amount of protein to perform the described experiments. The gene was synthesized and cloned into a pET28b vector, in frame with the N-terminal tag. *E. coli* cells of the strain BL21-Gold (DE3) were transformed, and protein was expressed in 1L of TB media with 50 mM kanamycin and 1mM of IPTG, by incubating overnight at 18 °C. Cells were harvested by centrifugation at 5129 g for 10 min, and pellets were resuspended in a solution of potassium phosphate 20 mM, at pH 7.5, with 500 mM NaCl, 25 mM imidazole, 1 mM PMSF and 0.1 mg/mL DNAse. The sample was sonicated and centrifuged at 20483 g for 20 min, afterwards, the supernatant was collected, filtered and loaded into a HisTrap^TM^ HP affinity column (GE Healthcare Bio-Sciences, Pittsburgh, PA, USA). The protein was eluted using a 0-500 mM imidazole linear gradient.

### Thermal-induced unfolding experiments by spectroscopies techniques

Thermal-induced unfolding was measured by fluorescence intensity and circular dichroism for LAO and dA, while only the second technique was used for dB. Samples of protein were prepared at 0.2 mg/mL in a buffer of bis-tris propane 10 mM for pH conditions above 7.0 and potassium acetates 10 mM for pH 5.0; these buffers were used for every experiment reported in this work. Fluorescence measurements were carried out on a PC1 ISS Spectrofluorometer (Champaign, IL-USA) equipped with a Peltier and a water-jacketed cell holder for temperature control, using a quartz cuvette of 1.0 cm pathlength. Tryptophan residues were excited with light at 295 nm and emission spectra were collected from 310 nm to 410 nm. CD measurements were carried out in a Chirascan VX spectropolarimeter (Applied Photophysics Leatherhead, England) using a quartz cuvette of 0.1 cm pathlength; ellipticity was recorded at 222 nm. Temperature was increased at a scan rate of 1.0 °C/min and data was collected every 0.5 min, from 25 °C to 75 °C.

The thermal-induced unfolding curves were normalized as protein unfolding fractions (#x1D453;_#x1D448;_) according to the next equation:

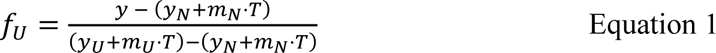

Where y is the detected signal, yN is the signal corresponding to the native state and mN its dependence on temperature, while yU is the signal of the unfolded state and mU its dependence on temperature, respectively.

The Van ’t Hoff enthalpy (ΔH_vH_) and melting temperature (T_m_) values were obtained by fitting the normalized curves to the next two-state equation, where R is the gas constant:

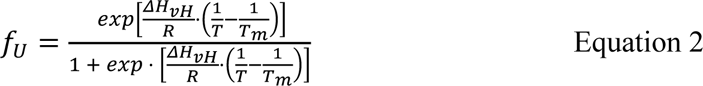

### DSC experiments

DSC experiments were carried out in a VP-Capillary DSC system (MicroCal, Malvern United Kingdom). Protein samples were dialyzed in the buffers at the desired pH conditions and prepared at a final concentration of 1mg/mL. Buffer baselines were acquired before the protein scans, using the same solutions utilized to dialyze the samples, and then subtracted to protein-buffer scans. Experiments were performed at a scan rate of 1.0 °C per minute, from 10 °C to 75 °C; after reaching the highest temperature, the samples were cooled down to the starting temperature and a second scan was initiated to measure the unfolding reversibility. Data were processed using the software Origin v.9.0 (OriginLab Corporation, Northampton, MA, USA.) with the MicroCal software extension for calorimetric data analysis. Baseline-subtracted scans were fitted to the next equation:

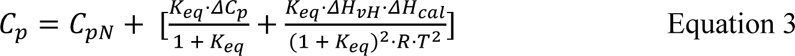

Where C_p_ is the molar heat capacity of the system, C_pN_ is the molar heat capacity of the folded state, K_eq_ is the equilibrium constant, and ΔH_vH_ the calorimetric enthalpy change. When a two-state model is assumed, ΔH_vH_ and ΔH_cal_ are the same.

### Equilibrium urea-induced unfolding experiments

The urea-induced unfolding of LAO and the individual domain A were measured by circular dichroism and fluorescence intensity; only CD was used to follow the unfolding on dB. Protein samples at 0.2 mg/mL were incubated overnight at 4 °C in urea concentrations ranging from 0 M to 5 M. Fluorescent measurements were obtained in an ISS PC1 spectrofluorometer (Champaign, IL-USA) as described above, using a quartz cuvette of 1.0 cm pathlength. Tryptophan residues were excited with light at 295 nm and emission spectra were collected from 310 to 410 nm at 25 °C. CD measurements were carried out in a Chirascan VX spectropolarimeter (Applied Photophysics, Leatherhead, England), using a quartz cuvette of 0.1 cm pathlength; signal was detected from 200 nM to 250 nm at 25 °C.

Urea-induced unfolding curves were normalized as protein unfolding fractions (#x1D453;_#x1D448;_) according to equation 1. In order to obtain the free energy change value in absence of denaturant (ΔG) and its dependence on the urea concentration (m), normalized curves were fitted to a simplification of the Santoro-Bolem equation [65]:

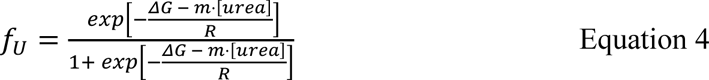

### Unfolding and refolding kinetics

Unfolding and refolding kinetics were carried out using a Chirascan VX spectropolarimeter (Applied Photophysics, Leatherhead, England) coupled to a stopped-flow module SX20. Experiments were measured on LAO and dA by circular dichroism and fluorescence intensity at pH 9.0; dB kinetics were acquired only by CD at pH 5.0. Changes in CD signal were monitored at 222 nm (230 nm for dB) while fluorescence intensity measurements were recorded using a 305 nm emission filter after excitation at 295 nm. Protein unfolding was performed by rapid mixing of protein samples into a buffer with the amount of urea necessary to reach final denaturant concentrations ranging from 3 M to 5 M. Refolding assays were performed by rapid mixing of protein samples, equilibrated overnight in 5 M of urea 10 mM bis-tris propane, into the buffer with the amount of urea required to reach final denaturant concentrations ranging from 2.75 M to 0.5 M. All the assays were carried out at final protein concentrations of 0.1 mg/mL, 25 °C.

### Analysis of Chevron plots

Chevron plots were analyzed in terms of a tree state model with an on-pathway intermediate (Scheme 1), assuming a rapid pre-equilibrium between I and U, according to the following expression [51]:

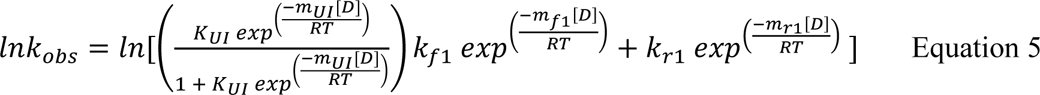

Chevron plots were also fitted to an off-pathway model (Scheme 2) according to the following expression:

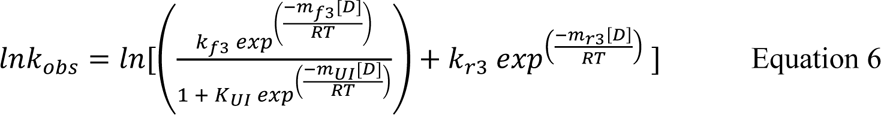

### Interrupted unfolding and refolding experiments

Interrupted unfolding (double jump) and refolding experiments were carried out on LAO at pH 9.0, 25 °C, using a Chirascan VX spectropolarimeter (Applied Photophysics Leatherhead, England) coupled to a stopped-flow module SX20.

For double jump assays, protein samples were unfolded by rapid mixing with urea 5 M for varying times (1 s-100 s) and subsequently refolded by diluting in buffer to reach a final urea concentration of 0.45 M. Refolding kinetics were monitored by circular dichroism at 222 nm and fitted to one or more exponentials to determine the number and magnitude of the rate constants after each unfolding time.

For interrupted refolding assays, protein was unfolded overnight at 4 °C in urea 5 M. Unfolded samples were refolded by rapid dilution with buffer and maintained in native conditions for varying times (0.5 s-30 s); native protein was subsequently unfolded by rapid mixing with urea to a final concentration of 5 M. Unfolding kinetics were monitored by CD at 222 nm and the rate constants were determined by fitting the curves to a two exponential function. The amplitudes associated with the rate constant were plotted versus refolding time. All the assays were carried out at final protein concentrations of 0.1 mg/mL and data was analyzed using the software Origin v.9.0 (OriginLab Corporation, Northampton, MA, USA.).

### Interrupted refolding analysis

The amplitude vs time plots obtained from the interrupted refolding experiments were fitted to the set of differential equations describing mechanisms 1-6, using the program R [66]. Schemes 1–3 are the simplest mechanisms which present only one intermediate (I), where U and N account for the unfolded and native states.

**Schema 1.**
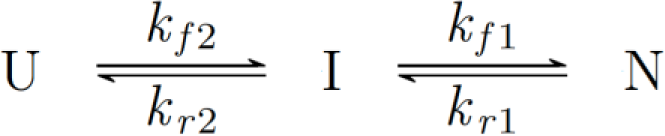

**Schema 2.**
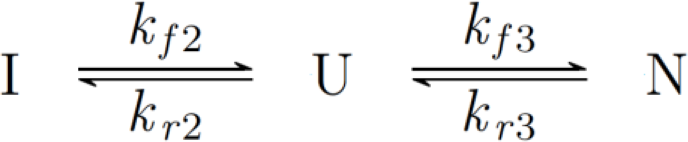

**Schema 3.**
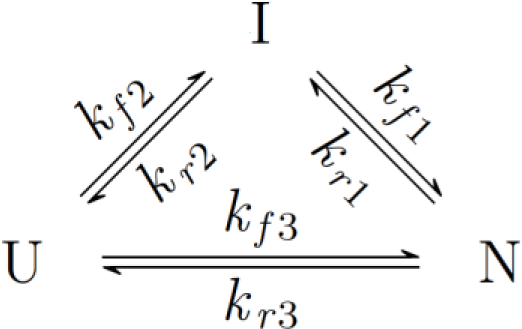

Schemes 4 and 5 represent folding mechanisms with two intermediates (I_C_ and I_T_). Scheme 5 is a branched system with a dead-end intermediate derived from scheme 4 when *k_f4_* and *k_r4_* are negligible, and U and I_T_ cannot interconvert directly, or do so at rates too low to have a detectable effect on the kinetics.

**Schema 4.**
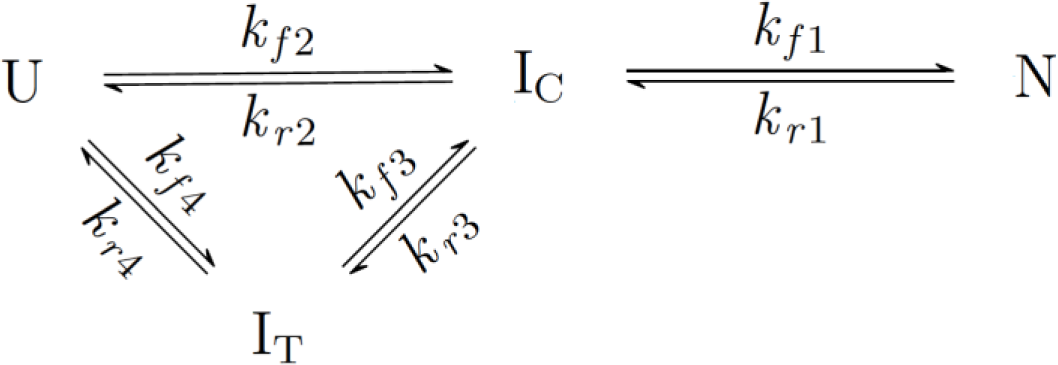

**Schema 5.**
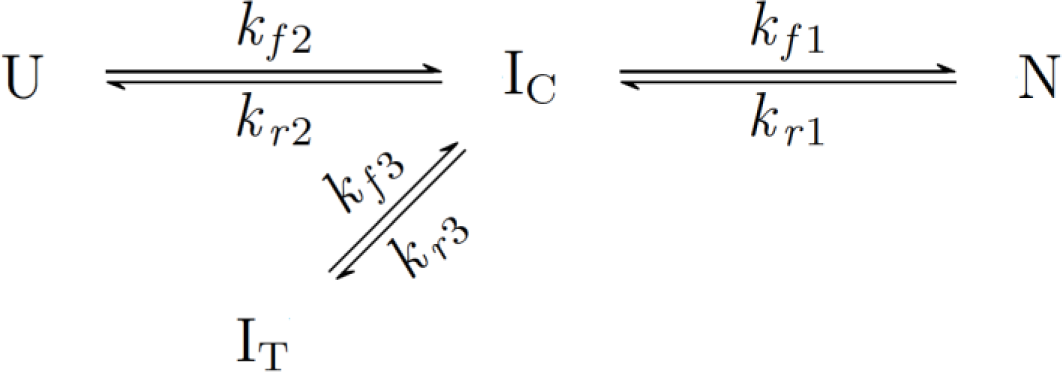

A more complex mechanism with four intermediate states is shown in scheme 6.

**Schema 6.**
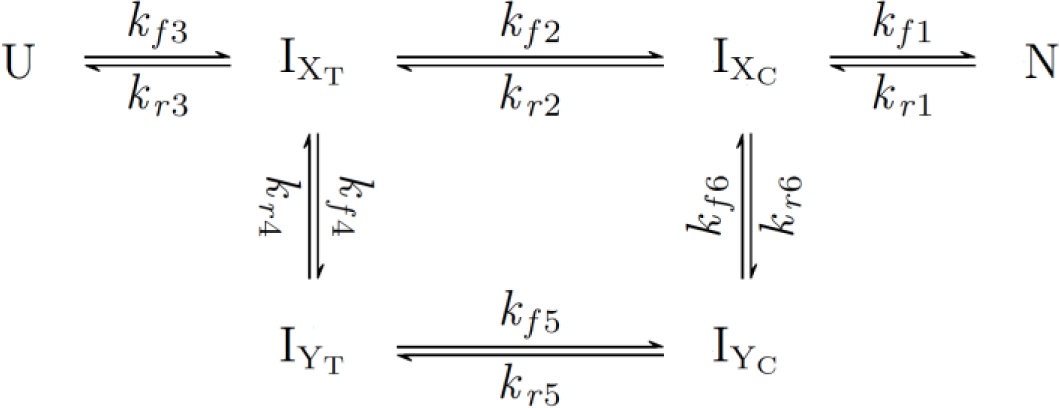

### Isothermal titration calorimetry experiments

ITC experiments were carried out on dA using a high-precision ITC200 microcalorimeter (GE Healthcare, Chicago, IL, USA). Arginine binding was measured at 25 °C in HEPES 10 mM, pH = 8.0. Protein was dialyzed exhaustively against the working buffer and loaded into the cell at a concentration of 0.27 mM. Arginine was dissolved in the same dialysis buffer to minimize dilution heats and loaded in the syringe at a concentration of 20 mM. 1uL injections of ligand were titrated in 200 s intervals until saturation was reached. Data was analyzed in the software Origin v.9.0 (OriginLab Corporation, Northampton, MA, USA.) with the MicroCal software extension for calorimetric data analysis.

The dissociation constant (K_d_), the enthalpy change (ΔH_b_), and the stoichiometry (n) were determined by nonlinear fitting of normalized titration data using a 1:1 binding model.

### Crystallization and three-dimensional structure determination of dA

For crystallization, domain A from LAO was exhaustively dialyzed in HEPES 10 mM pH: 8.0 after concentrated with Amicon Ultra centrifugal filter units (Millipore) at a final concentration of 13.5 mg mL^−1^. Crystallization plates using a hanging-drop vapor diffusion method were set up with a final volume of 3.0 µL and 1:1 protein: mother liquid ratio. Diffracting crystals grew at 18 °C after 6 weeks in 0.2 M sodium acetate trihydrate, 0.1 M sodium cacodylate trihydrate pH: 6.5, 30% w/v polyethylene glycol 8000. Diffraction data were collected at 100 K at X-ray in-house source (wavelength: 1.5418 Å) generated by a Rigaku MicroMax-007 HF rotating anode (Rigaku, The Woodlands) using an oscillation of 0.5 degrees by frame on a DECTRIS PILATUS 200K image plate detector (DECTRIS, Baden-Daettwil). Datasets were processed with HKL-3000 [67]. The number of molecules per asymmetric unit (ASU) was determined by calculating Matthew’s coefficient, yielding a value of 2.28 corresponding to 3 copies by ASU. Molecular replacement was performed with PHASER in the PHENIX software suite v.1.19.2 [68] using the edited pdb file corresponding to the domain A from *Salmonella typhimurium* LAO [43] (PDB ID: 6MLE) as a starting model. Data refinement was performed with phenix.refine [69] and iterative manual model improvement by rebuilding in COOT v.0.9 [70]. The final tridimensional structure was validated with MolProbity [71] and Protein Data Bank validation server, satisfying all quality criteria. Coordinates and structure factors were deposited in the PDB database https://www.rcsb.org/ with the accession code 6XKS. Table S2 summarizes data collection and refinement statistics. The figures were created using PyMOL Molecular Graphics System v.4.6.0 (Schrodinger, LLC).

## Author contributions

R.V., D.A.F-V., D-A.S. and A.S-P designed the study; R.V., T.B. and E.I.J.M performed the temperature and urea-induced unfolding assays, the unfolding and refolding kinetics, and data analysis; R.V. carried out and analyzed the interrupted refolding and double jump assays; S.R-R and A.R-R. performed x-ray crystallography; I.V-L. was in charge of protein expression and purification; N.O.P. performed ITC experiments; H.A.L.S, expressed and purified cyclophilin A, and participated in the design and analysis of the kinetic studies, D-A.S. designed the isolated domain constructs; R.V., T.B, E.I.J.M., S.R.R., and M.C. performed and analyzed DSC experiments; R.V., D.A.F-V. and R. R-S. evaluated the kinetic data to provide mechanistic information of the folding process; A.S-P. and D.A.F-V. supervised research and provided financial support; R.V. and D.A.F.-V. wrote the manuscript. All the authors revised the manuscript.

## Acknowledgements

This work was supported by Programa de Apoyo a Proyectos de Investigación e Innovación Tecnológica DGAPA-UNAM (PAPIIT grant number IN219519 to DAFV), Facultad de Medicina, UNAM. R.V., T.B., E.I.J.M. and H.A.L.S. received CONACYT scholarships for graduate studies. X-ray structures were determined at the Laboratorio Nacional de Estructura de Biomacromoléculas (LANEM).

## Notes

### Competing Interest Statement

The authors have declared no competing interest.

