## Supplementary Information for "Thermodynamic and kinetic analysis of the LAO binding protein and its isolated domains reveal non-additivity in stability, folding, and function"

### SUPPORTING INFORMATION

#### Table S1. Rate constants from interrupted unfolding assays

| | Scheme 1 | Scheme 2 | Scheme 3 | Scheme 4 | Scheme 5 | Scheme 6 |
|---|---|---|---|---|---|---|
| $k_{f1}$ | 0.17 ± 0.01 | | 0.12 ± 0.01 | 3.2 ± 0.57 | 4.1 ± 0.3 | 5.1 ± 3.2 |
| $k_{r1}$ | $1.8 \times 10^{-5} \pm 5 \times 10^{-6}$ | | $3.0 \times 10^{-4} \pm 2 \times 10^{-3}$ | $1.2 \times 10^{-4} \pm 2 \times 10^{-5}$ | $1.0 \times 10^{-4} \pm 8 \times 10^{-6}$ | $9.3 \times 10^{-3} \pm 1.6 \times 10^{-4}$ |
| $k_{f2}$ | 0.6 ± 0.06 | 0.7 ± 0.14 | 0.45 ± 0.05 | 0.58 ± 0.02 | 0.59 ± 0.02 | 1.0 ± 0.6 |
| $k_{r2}$ | 0.11 ± 0.04 | 0.38 ± 0.08 | 0.03 ± 0.03 | 1.1 ± 0.25 | 1.0 ± 0.07 | 3.9 ± 2.5 |
| $k_{f3}$ | | 0.29 ± 0.04 | 0.08 ± 0.01 | 0.8 ± 0.12 | 0.8 ± 0.08 | 0.58 ± 0.06 |
| $k_{r3}$ | | $5.5 \times 10^{-5} \pm 3 \times 10^{-3}$ | $3.0 \times 10^{-4} \pm 2 \times 10^{-3}$ | 16.3 ± 1.8 | 22.8 ± 1.9 | 0.21 ± 0.3 |
| $k_{f4}$ | | | | $4.6 \times 10^{-4} \pm 8 \times 10^{-5}$ | | 2.3 ± 0.8 |
| $k_{r4}$ | | | | $9.5 \times 10^{-4} \pm 1 \times 10^{-4}$ | | 0.9 ± 0.4 |
| $k_{f5}$ | | | | | | 0.08 ± 0.06 |
| $k_{r5}$ | | | | | | 8.4 ± 8.4 |
| $k_{f6}$ | | | | | | 14.5 ± 9.5 |
| $k_{r6}$ | | | | | | 1.6 ± 1.3 |

#### Table S2. Data collection and refinement statistic for unligated dA

| Data collection[a] | |
|---|---|
| PDB ID | 6XKS |
| Wavelength (Å) | 1.5418 |
| Crystallization conditions | 0.2 M Sodium acetate trihydrate<br>0.1 M Sodium cacodylate trihydrate pH: 6.5<br>30% w/v Polyethylene glycol 8000. |
| Resolution range (Å) | 42.86 – 2.40 (2.44 – 2.40) |
| Space group | P 21 21 21 (19) |
| Unit cell dimensions | |
| a, b, c, (Å) | 46.60, 87.78, 109.15 |
| α, β, γ, (°) | 90.0, 90, 90.0 |
| Total reflections | 220497 (7948) |
| Unique reflections | 32910 (1691) |
| Multiplicity | 6.7 (4.7) |
| Completeness (%) | 99.3 (98.6) |
| Mean I / sigma (I) | 16.6 (2.0) |
| Wilson B-factor | 31.4 |
| R-merge | 0.160 (0.826) |
| CC1/2 | 0.991 (0.694) |
| CC* | 1.000 (0.899) |
| Matthew's coefficient Vm (Å3 Da-1) | 2.28 |
| Solvent content (%) | 46.1 |
| Protein molecules per asymmetric unit | 3 |
| **Refinement statistics** | |
| Reflections used in refinement | 17859 (1691) |
| Reflections used for R-free | 1785 (169) |
| Rwork , Rfree | 0.232 (0.295), 0.285 (0.366) |
| Number of atoms | 3460 |
| Protein | 3282 |
| Ligand | 55 |
| Water | 152 |
| Protein residues | 432 |
| RMS (bonds) (Å) | 0.005 |
| RMS (angles) (°) | 0.730 |
| Ramachandran favored (%) | 96.01 |
| Ramachandran allowed (%) | 3.99 |
| Ramachandran outliers (%) | 0.00 |
| Rotamer outliers (%) | 0.30 |
| Clashscore | 4.82 |
| Average B-factor | 56.3 |
| Protein | 56.0 |
| Ligand | 68.9 |
| Solvent | 58.7 |
| TLS groups | 3 |

[a] Statistics for the highest-resolution shell are shown in parentheses.

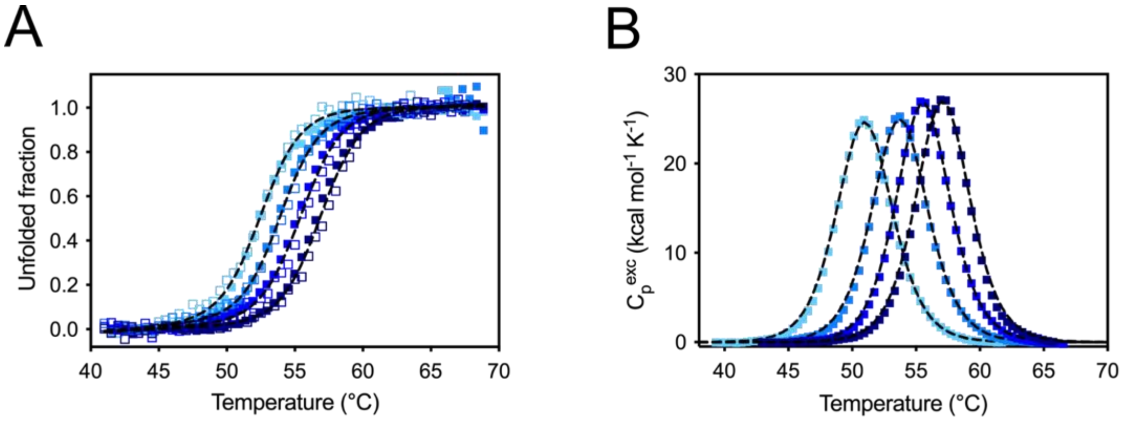


**Figure S1.** Thermal unfolding of LAO measured by spectroscopic techniques (CD as open and IF as closed symbols) (A) and DSC (B) at pH 8.0 (sky blue), 8.5 (azure blue), 9.0 (royal blue) and 9.5 (navy blue). Data were fitted to a two-state model (black lines).

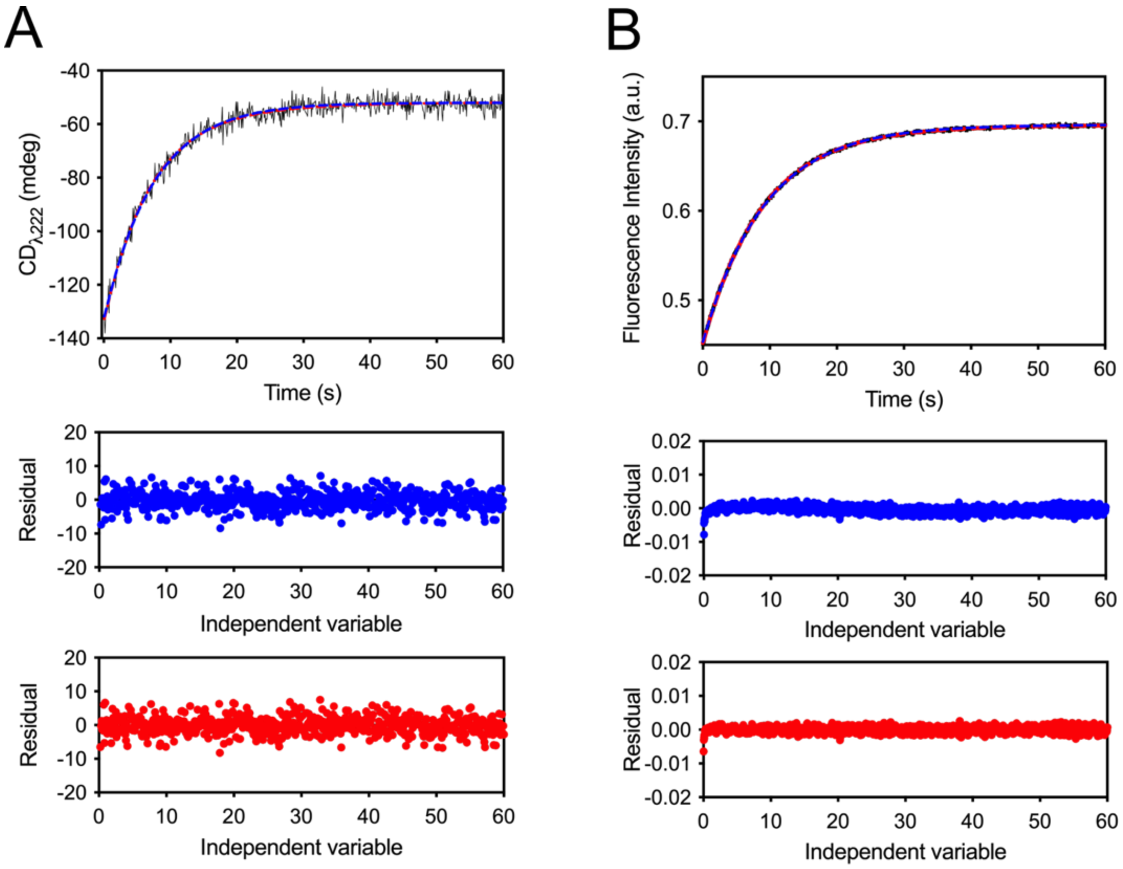


**Figure S2.** Unfolding kinetics of LAO measured by CD (A) and IF (B); a final urea concentration of 5.0 M was reached. Kinetic traces were fitted to one (blue dashed lines) and two (red dashed lines) exponential functions; the corresponding residual plots (below) are shown as dots of the same color as the fitting curves. Both CD and IF data were well described by one exponential with a rate constant of 0.15 $s^{-1}$.

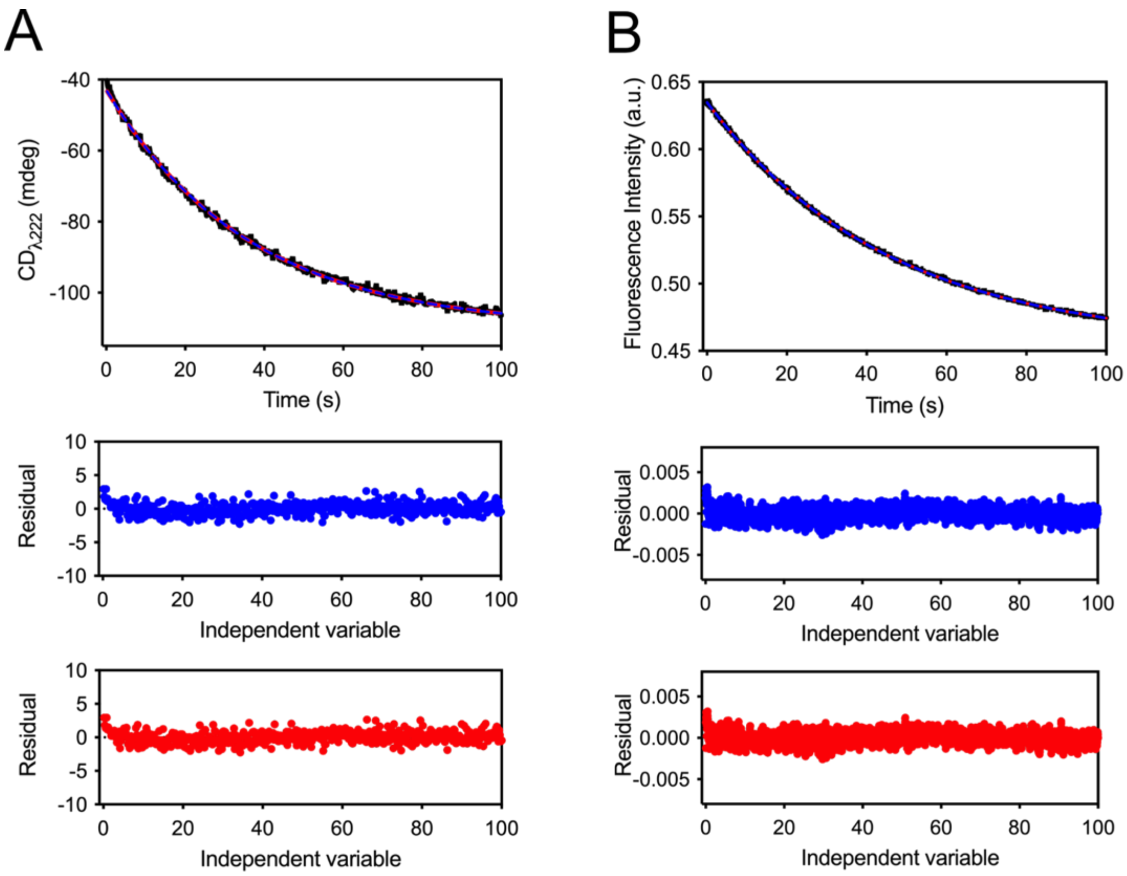


**Figure S3.** Refolding kinetics of LAO measured by CD (A) and IF (B); a final urea concentration of 2.0 M was reached. Kinetics traces were fitted to one (blue dashed lines) and two (red dashed lines) exponential functions; the corresponding residual plots (below) are shown as dots of the same color as the fitting curves. Both CD and IF data were well described by one exponential with a rate constant of 0.02 $s^{-1}$.

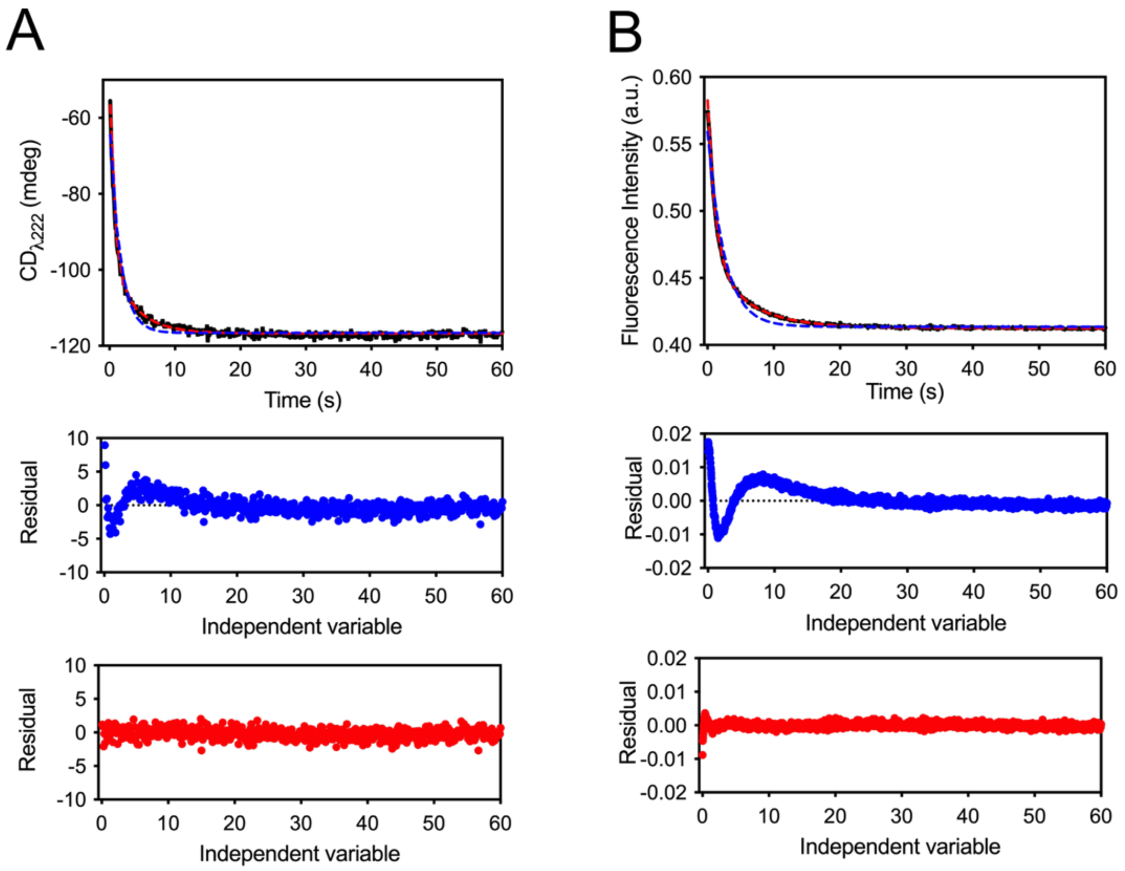


**Figure S4.** Refolding kinetics of LAO measured by CD (A) and IF (B); a final urea concentration of 0.45 M was reached. LAO was previously denatured in 5M urea for 24h. Kinetics traces were fitted to one (blue dashed lines) and two (red dashed lines) exponential functions; the corresponding residual plots (below) are shown as dots of the same color as the fitting curves. Both CD and IF data were best described by two exponentials with rate constants of 0.9 $s^{-1}$ and 0.18 $s^{-1}$.

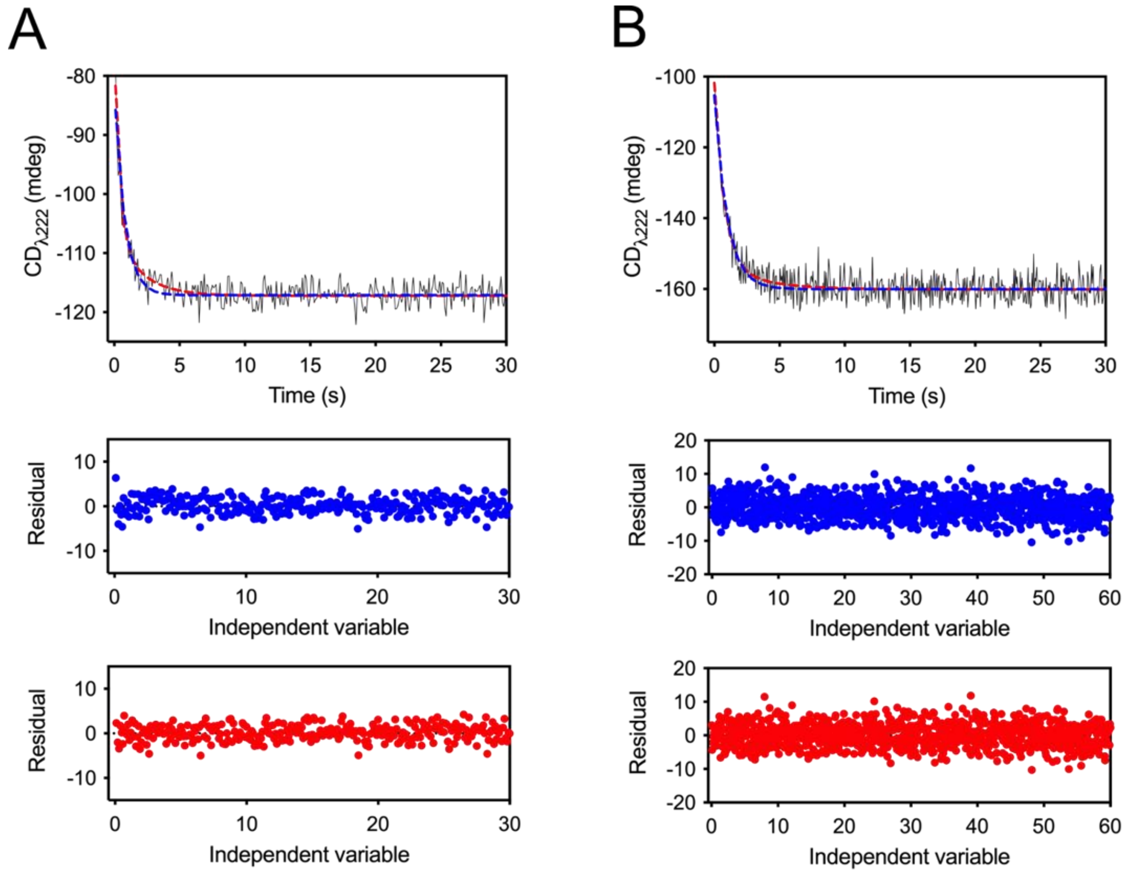


**Figure S5.** A) Refolding kinetics of LAO, at a final urea concentration of 0.45 M, measured by CD in the second step of the double jump experiment after 3 s delay. Kinetic traces were fitted to one (blue dashed lines) and two (red dashed lines) exponential functions. B) Kinetic data acquired from simple-mixing experiments in the presence of 5 uM of cyclophilin A were well described by one exponential function with a rate constant of 1.0 $s^{-1}$. Residual plots (below) are shown as dots in the same color as its corresponding fitting curves.

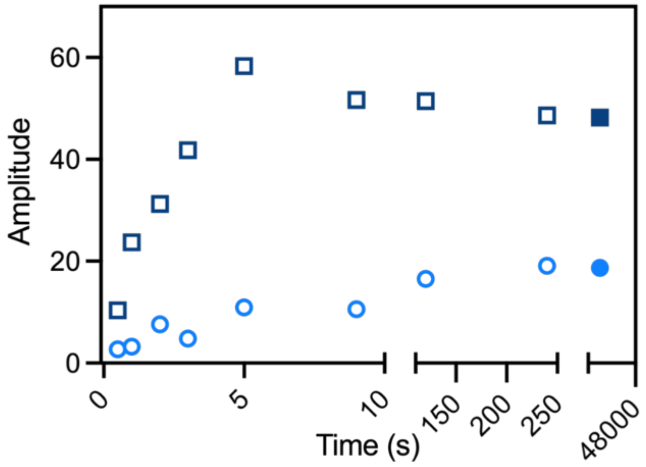


**Figure S6.** Amplitudes associated with the fast (open squares) and slow (open circles) constants obtained from the double jump refolding kinetics. Amplitudes from the refolding kinetics are shown in Figure S4A are shown as closed symbols.

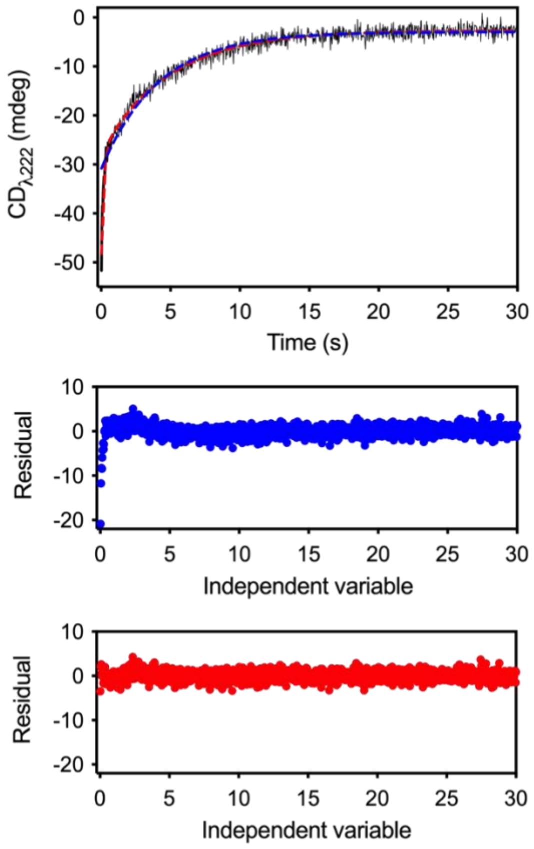


**Figure S7.** Unfolding kinetics of LAO measured by CD in the second step of the interrupted refolding experiment after 5 s delay; a final urea concentration of 5 M was reached. Kinetic traces were fitted to one (blue dashed lines) and two (red dashed lines) exponential functions; the corresponding residual plots (below) are shown as dots of the same color as the fitting curves. Data obtained at all delay times were well described by two exponentials with rate constants of 0.2 $s^{-1}$ and 6.3 $s^{-1}$.

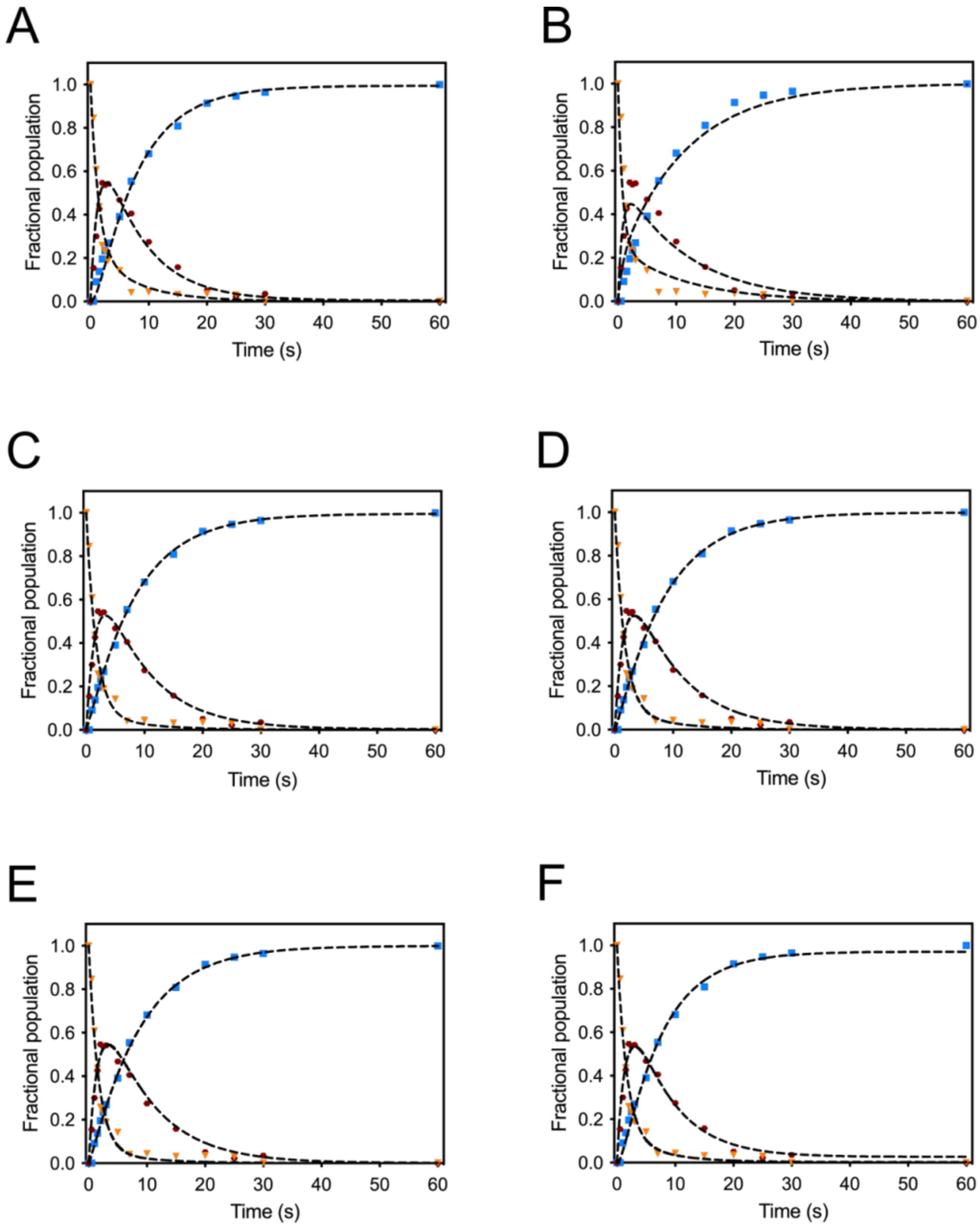


**Figure S8.** Time evolution of the protein states populated by LAO during refolding. The amplitudes associated with the slow rate constant obtained from the interrupted refolding kinetics report on the fraction of molecules in the native state (blue dots), whereas the amplitude of the fast rate constant (Figure S7) was ascribed to the fraction of molecules in one or more intermediate states (red dots). Fractional populations were normalized considering as 1.0 the amplitude of the 60 s delay kinetic, where only the slow phase was observed. The fraction of unfolded molecules (orange dots) was obtained by difference at each delay time. Data were fitted to the schemes 1 to 5 (shown in panels A to B, respectively).

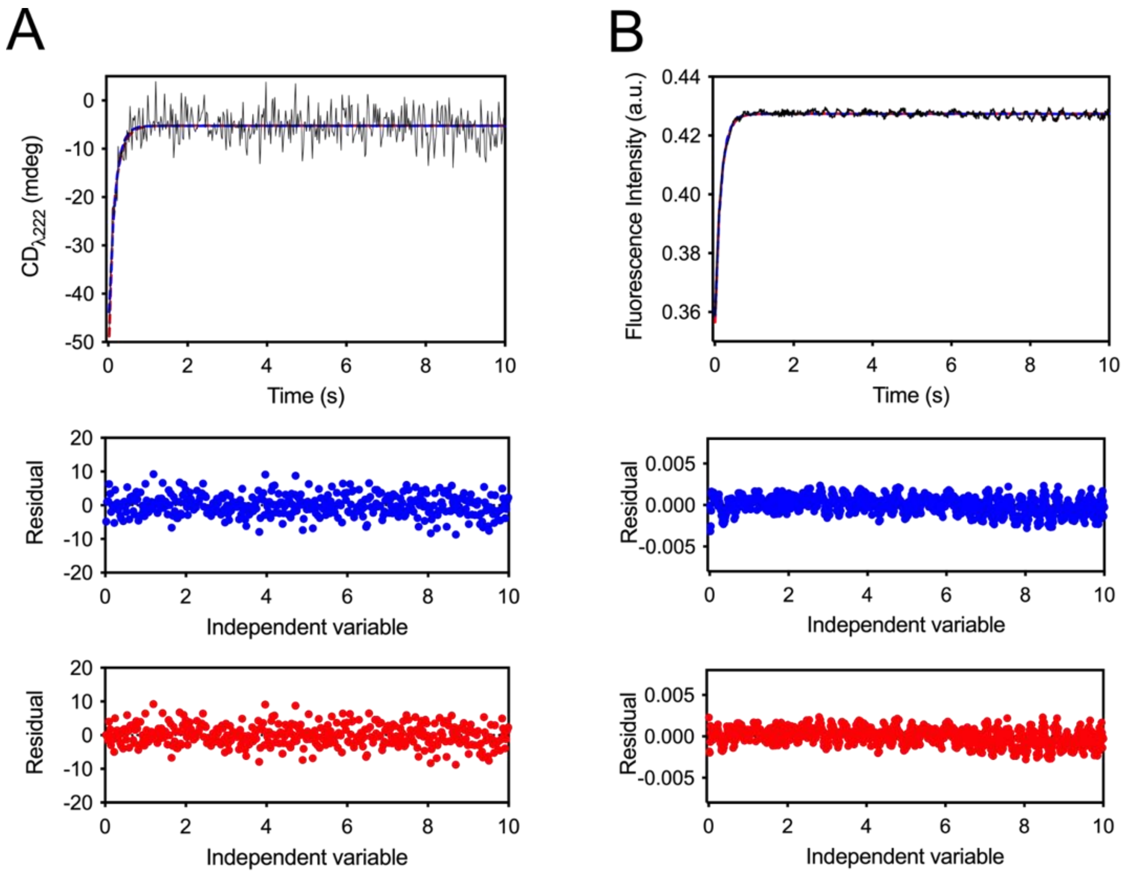


**Figure S9.** Unfolding kinetics of dA measured by CD (A) and IF (B); a final urea concentration of 5 M was reached. Kinetics were fitted to one (blue dashed lines) and two (red dashed lines) exponential functions; the corresponding residual plots (below) are shown as dots of the same color as the fitting curves. Data was well described by one exponential with a rate constant of 6.7 $s^{-1}$.

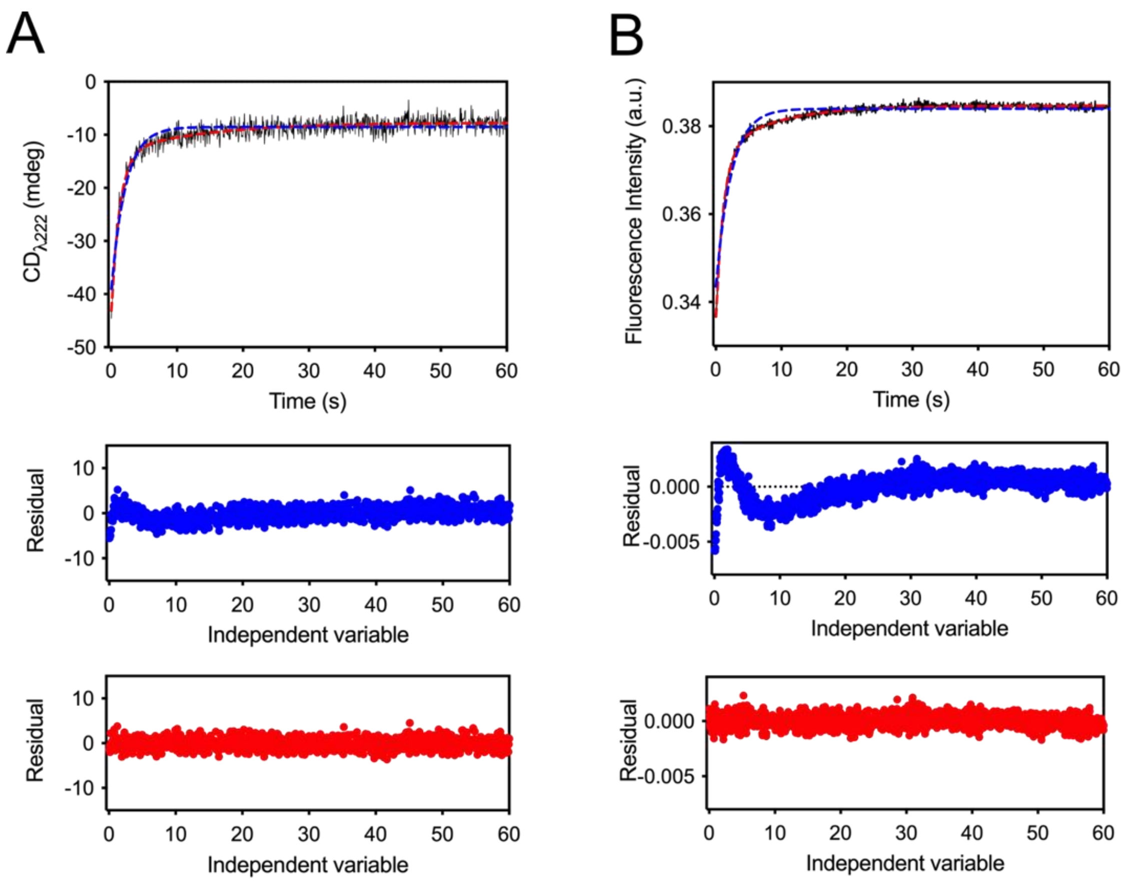


**Figure S10.** Unfolding kinetics of dA measured by CD (A) and IF (B); a final urea concentration of 3 M was reached. Kinetics were fitted to one (blue dashed lines) and two (red dashed lines) exponential functions; the corresponding residual plots (below) are shown as dots of the same color as the fitting curves. Data was best described by two exponentials with rate constants of 0.06 $s^{-1}$ and 0.7 $s^{-1}$.

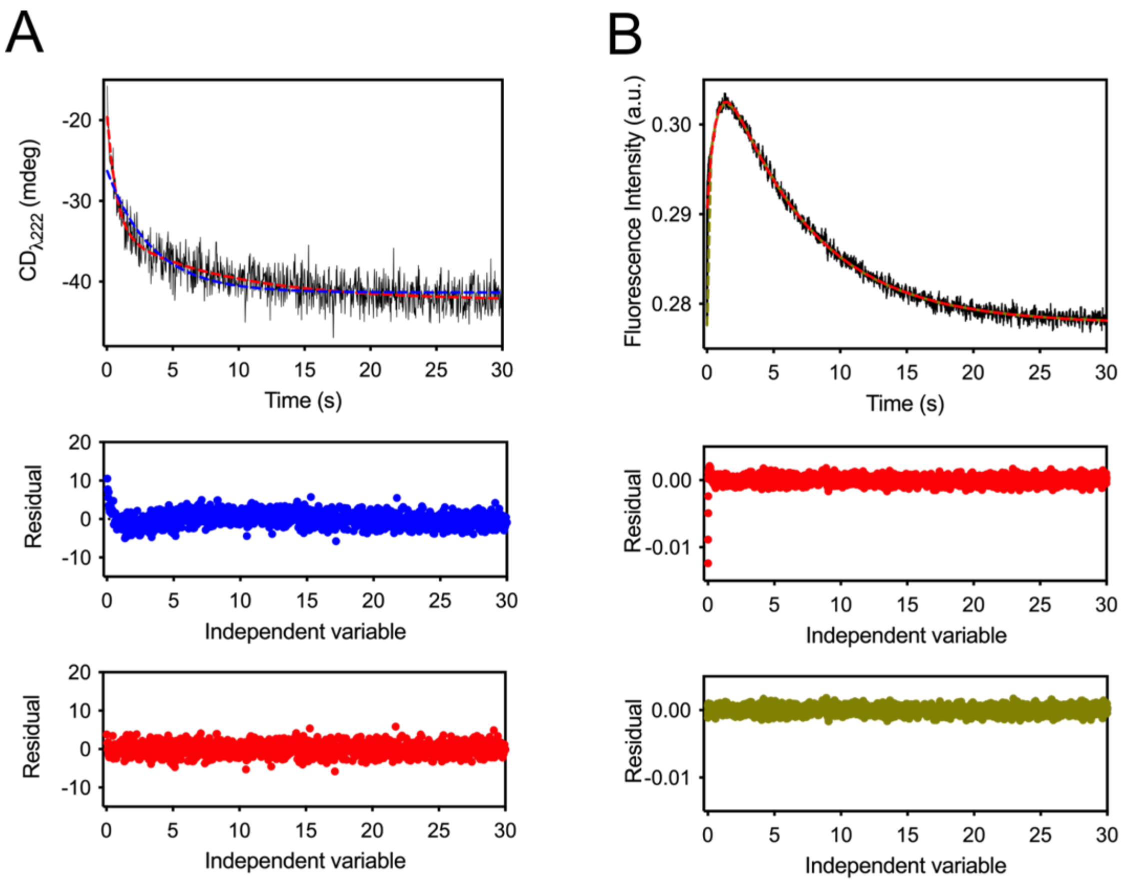


**Figure S11.** Refolding kinetics of dA measured by CD (A) and IF (B); a final urea concentration of 0.45 M was reached. Kinetics were fitted to one (only CD, blue dashed lines) , two (CD and IF, red dashed lines) and three (only IF, green dashed lines) exponential functions; the corresponding residual plots (below) are shown as dots of the same color than the fitting curves. CD data was best described by two exponentials with rate constants of 0.16 $s^{-1}$ and 1.5 $s^{-1}$, while IF data were best described by three exponentials with rate constants of 0.16 $s^{-1}$, 1.3 $s^{-1}$ and 61.4 $s^{-1}$.

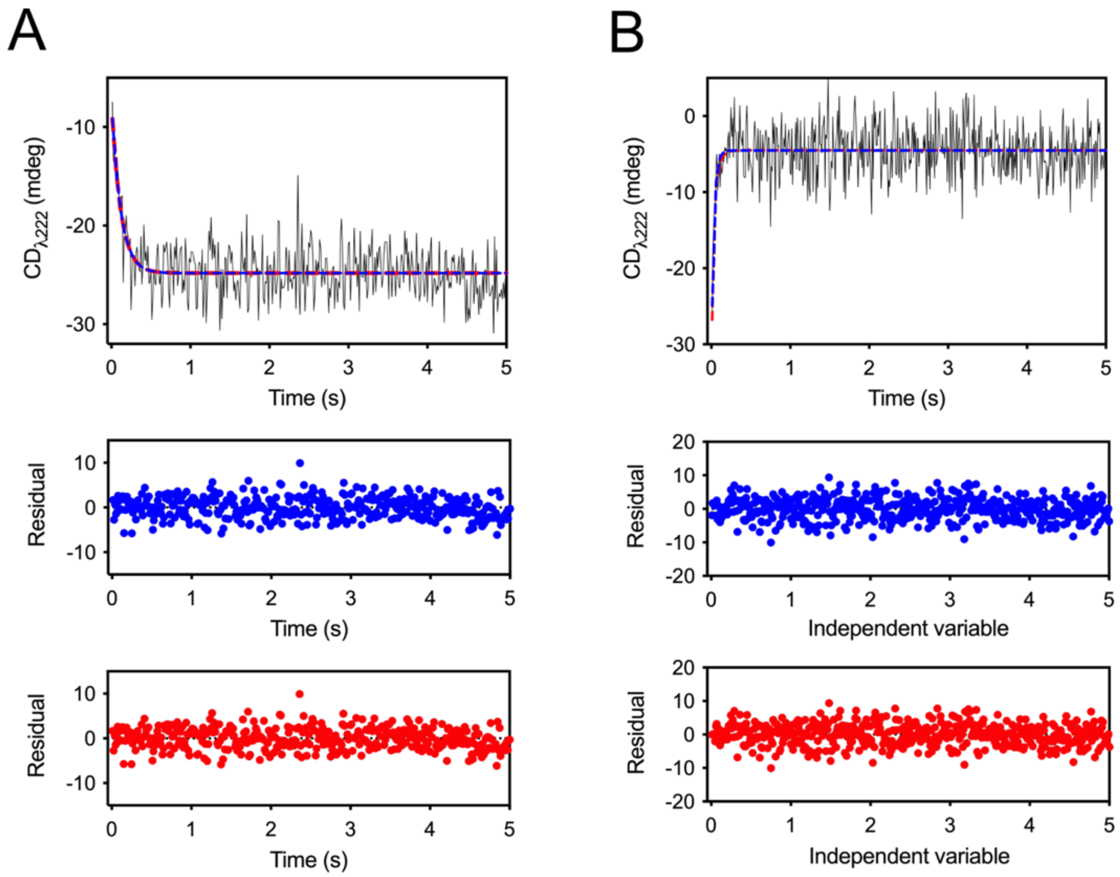


**Figure S12.** Refolding (A) and unfolding (B) kinetics of dB measured by CD; final urea concentrations of 0.45 M and 3 M were reached in refolding and unfolding, respectively. Kinetics were fitted to one (blue dashed lines) and two (red dashed lines) exponential functions; the corresponding residual plots are shown as dots of the same color as the fitting curves. Data were well described by one exponential with a rate constant of 14.5 $s^{-1}$ for unfolding and 10.9 $s^{-1}$ for refolding.
